# The distance decay of similarity in tropical rainforests. A spatial point processes analytical formulation

**DOI:** 10.1101/216929

**Authors:** Anna Tovo, Marco Favretti

## Abstract

In this paper we are concerned with the analytical description of the change in floristic composition (species turnover) with the distance between two plots of a tropical rainforest due to the clustering of the individuals of the different species. We describe the plant arrangement by a superposition of spatial point processes and in this framework we introduce an analytical function which represents the average spatial density of the Sørensen similarity between two infinitesimal plots at distance *r*. We see that the decay in similarity with the distance is essentially described by the pair correlation function of the superposed process and that it is governed by the most abundant species. We test our analytical model with empirical data obtained for the Barro Colorado Island and Pasoh rainforests. To this end we adopt the statistical estimator for the pair correlation function in [1] and we design a novel one for the Sørensen similarity. Furthermore, we test our analytical formula by modelling the forest study area with Neyman-Scott point processes. We conclude comparing the advantages of our approach with other ones existing in literature.

## Introduction

Estimating biodiversity of forests is a central issue in modern conservation ecology. Both from the theoretical and field application point of view it represents a daunting challenge. Since the pioneering work of Whittaker [2] and Preston [3, 4] a number of diversity indices have been introduced in literature and their effectiveness has been tested against field data, with various degrees of success. In this paper we are concerned with a single aspect of this broad issue, namely the study of the decay of similarity between two regions of a landscape as a function of the distance between them. To specify the intuitive concept of similarity we will adopt the widely used Sørensen^1^ similarity index [5, 6] (see equation (2) below) and its associated spatial density (equation (3)). Equally used in literature is the notion, complementary to the concept of similarity, of species turnover or β-diversity, that is the change in species composition between two plots as a function of the distance between them. Even stated in these terms, this more restricted problem is hard to reduce to a mathematical model since on real landscapes many drivers of diversity are acting at the same time and may contribute with different intensity depending on the spatial scales [7]: at a continental scale climatic factor may dominate whereas at a smaller scale orographic factors may create specific environmental gradients due to the change in altitude or to the orientation of valleys. At any scale, the effect of past transformations of the environment may have shaped the territory with dispersal barriers or niches. The heterogeneity of these factors may have hampered the construction of a all-compassing mathematical model and, effectively, a relatively small (compared the huge number of articles dedicated to biodiversity issues) number of works are available on the specific problem of finding the function that best describes the change in species composition with the distance. In chronological order, important contributions to this central problem of estimating biodiversity of forests are the seminal works of Leight et al. [8], Nekola and White [9] and the neutral theory approach of Hubbell [10, 11] (see e.g. the comprehensive book Magurran and McGill [12]).

In this paper we focus on a single driver of diversity, that is the tendency of plants to form clusters of individuals. The shape and extent of the cluster may vary from species to species depending on seed dispersal limiting factors, or other effects, e.g Janzen-Connell effect [13, 14], which may be inter o intra specific. We aim at reducing this multiplicity of biotic factors to a single statistical descriptor. Stated in more mathematical terms, we wish to study the effect of spatial correlations between the individuals’ (plants) positions on the change in the species’ composition of two small plots at a given distance. The mathematical tool adapted to this task is the spatial point process theory (see [15, 16]). The main sources of inspiration for our approach are the works of Shimatani [1, 17], Plotkin and al. [18], Morlon et al. [19] (based on [18]), and also Chave et al.[20]. Using the language of point processes we derive in Sect. 1 an analytical formula for χ(*r*), the (spatial density of) Sørensen similarity between two small plots distance *r* apart. This gives the form of the decay of similarity as a function of the distance:

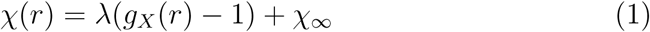

where the first term *g*_*X*_(·) is the pair correlation function of the superposed process having intensity λ and χ_∞_ is a constant depending solely on the species’ abundances and representing the similarity at a scale where the clustering of individuals has no effect. Thus, essentially, the decay in similarity is described by the pair correlation function of the whole forest plot. This latter in turn depends on the clustering of each species *weighted by their relative abundance* (see equation (15)). Therefore in our model the similarity decay function is dominated by the most abundant species, a feature previously recognized in other studies [19], but still debated [6, 21].

Apart from presenting a novel analytical approach to the definition of a decay of similarity function using point processes, the main aim of this paper is to test the proposed formula against field data. In our study we use the BCI (Barro Colorado Island) and Pasoh forest databases, which register the spatial position of respectively 222602 and 310520 plants belonging to 301 and 927 species covering an area of 50ha each. In Sect. 2 we introduce the statistical estimators for the similarity decay, 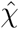, and for the pair correlation function, 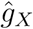 The former, based directly on Sørensen similarity formula (2), as far as we know is a novel one, whereas the latter has been used in [1] and it is derived from the general theory of point processes even if it *does not need any hypothesis on the type of stochastic point process* that we should associate to the species of the forest under study. They therefore provide a test of the formula (1) above at a very general level.

A second goal of this work is to select the class of spatial point process that best describes the plants’ arrangement in the study area and test its effectiveness in reproducing the decay of similarity function. This is done in Sect. 3. The clustering of each species α is described by a univariate (if we assume rotational symmetry of the two-dimensional cluster) probability density *d_α_*(*r*), the so-called dispersal kernel, which gives the probability that an individual of the cluster will establish at distance *r* from its parent, located at the cluster’s center. The dispersal kernel features of each species are thus the essential informations for our model that have to be drawn from experimental data (by the minimum contrast method in this work). We test the effectiveness of three dispersal kernels (exponential, Gaussian, Cauchy) at describing the species’ clustering. Once we determine the cluster parameters for each species, we compute the analytical form of the pair correlation function *g_X_* and of the similarity index χ(*r*). These are compared with their statistical estimates in Sect. 4.3.

## ^1.^ Similarity decay functions

### ^1.1^ Sørensen index for point processes

We begin by recalling the definition of Sørensen similarity index. We consider a flat region *W* with no environmental gradients. Given two disjoint (*A ∩ B* =) subregions *A* and *B*, let *s*(*A*) and *s*(*B*) be the number of different species present, respectively, in *A* and *B* and let *s*(*A, B*) be the number of co-present species in *A* and *B*. Provided that *s*(*A*) + *s*(*B*) *>* 0, the Sørensen similarity between regions *A* and *B* is the symmetric function 0 ≤ *σ*(*A, B*) ≤ 1

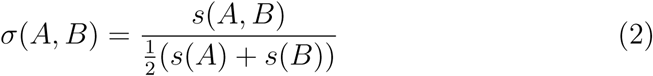

When *s*(*A*) = *s*(*B*), σ(*A, B*) gives the number of co-present species per species. As it is well known, the number of present or co-present species depends on the size of the regions *A* and *B*. Therefore we assume, as it is generally the case, that *A* and *B* have the same size *a* and we denote with

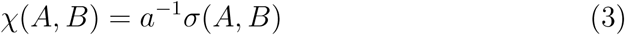

the number of co-present species per species and per unit of survey area, i.e. the spatial density of Sørensen similarity.

In the same spirit of [1], we wish now to reformulate the notion of Sørensen similarity in the language of spatial point processes. We model the presence of *S* species by a spatial point process 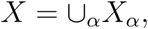 =∪_α_*X*_α_, which is the superposition of *S* mutually independent, homogeneous and isotropic spatial point processes *X*_*α*_, α *∈* {1, …, *S*}. In this way we model a community of *S* independent species where intra-specific interactions are allowed. Let us denote with *n*_*α*_(*x*) and *n*_*α*_(*y*) the random number of points of the process *X*_*α*_ contained in two infinitesimal disks centered at *x* and *y*, having equal area *dx* = *dy* and being disjoint. Therefore 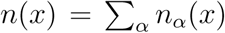 is the total number of individuals in *dx*, regardless of their species.

Let us now introduce some basic notions on point process theory. Let λ_α_(*x*) be the intensity (spatial density of points) of 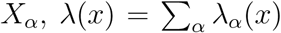 be the intensity of the superposed process *X*, ρ_α_(*x, y*) be the associated second order product density (second moment density, see [16] or [22]) and set for simplicity’ sake 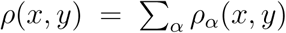 The following interpretations are standard

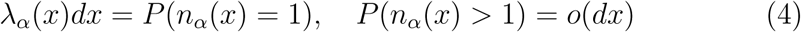

and

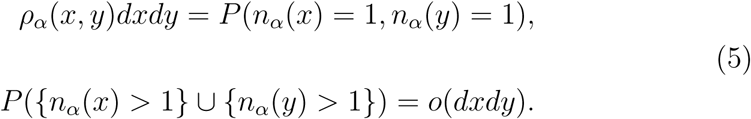

Denoting with *s*(*x*), *s*(*y*) and *s*(*x, y*) the number of species (point processes) present in *dx*, *dy* and in both *dx* and *dy* respectively, and neglecting higher order terms in *dx* or *dxdy*, we have that the *expected number* of species found in *dx* around *x* can be expressed as

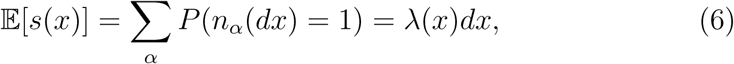

while the average number of co-present species in the infinitesimal regions *dx* and *dy* around *x* and *y* is

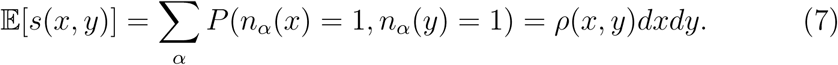

The number *s*(*x*) of species at *x* and the number *s*(*x, y*) of shared species at *x* and *y* are discrete random variables whose expected values can be described by the above formulae (6) and (7). Apart from their averages, their distribution is assumed to be unknown. The equivalent of the Sørensen similarity index for infinitesimal regions is thus given by the random variable

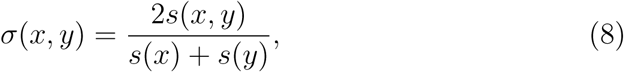

which is the ratio of two random quantities. The expected value of this ratio can be computed from 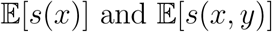 using the method of statistical differentials. Indeed, let us denote with *X* and *Y* the two random variables *s*(*x, y*) and *s*(*x*) + *s*(*y*), respectively. Then (see e.g. [23], p.65)

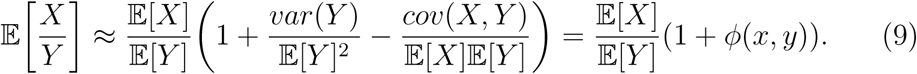

The point process formulation of the Sørensen similarity index can therefore be computed as

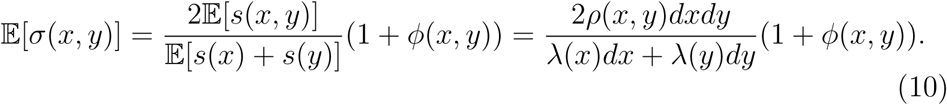

Considering the associated spatial density

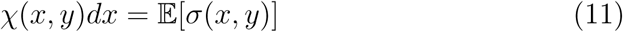

we have the following form for the similarity density

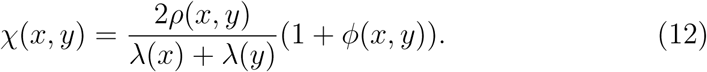

The factor 1 + *ϕ*(*x, y*) is hard to compute analytically. In Sect. 2 we show that for our applications it can be set equal to zero.

### 1.2. Similarity decay under φ(*x, y*) = 0, stationarity and isotropy hypotheses

Let us consider the special case where φ(*x, y*) equals 0 for all *x* and *y* in our study region *W*. If ρ_α_(*x, y*) is a function of the distance *r* = |*x – y*| (isotropy), and if the intensity (giving the expected density of species in our model, see (6)) is constant (stationarity), i.e. λ(*x*) = λ, formula (12) above becomes distance dependent

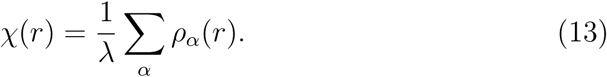

If we rewrite 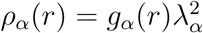 using the pair correlation function *g*_*α*_ and we and we introduce the relative intensity *p*_*α*_ = *λ*_*α*_/*λ*, with ∑_α_*p*_*α*_ = 1, equation (13) becomes

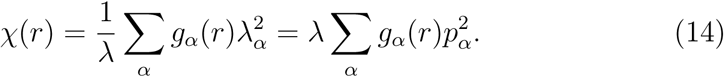

Introducing the pair correlation function *g*_*X*_(*r*) of the superposed process [16]

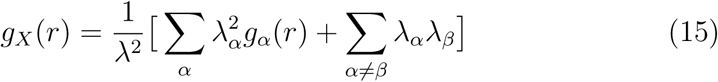

we obtain the Sørensen similarity decay function (13) in the form

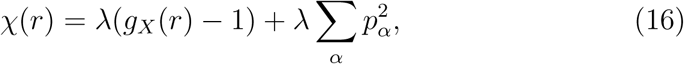

which coincides with the pair correlation function of the superposed process up to a change of scale.^2^ This is the main object of our paper.

Unlike the original Sørensen index (2) which is an incidence-based index, i.e. giving equal weight to abundant or rare species, this one takes into account the relative intensity of species *p*_*α*_. As noted in [6], incidence-based indices are generally biased downward, underestimating similarity especially when species richness is large or sample size is small. From formula (14) we notice that the similarity decay function we propose is dominated by the most abundant species, a feature which has been previously recognized in the literature (see [19]), but that is rigorously motivated in our formula.

A useful property of the formula (14) above is that it is independent of the size of the plots. This is the consequence of our strategy of considering virtually infinitesimal plots and considering the spatial density of Sørensen similarity. In a sense, our approach is orthogonal to the one adopted in the works [19] and [18] where the emphasis is first on the development of an analytical formula for the similarity between plots of finite area (even if the assumption of relatively small sample size and relatively large distances is made in [19]) and then on dealing with the problem of determining how the model parameters (e.g. the negative binomial clumping parameter in [18]) vary with the area under study. See Sect. 5 below for an overview on this problem. We think that our approach, even if not all-compassing, is more straightforward and easier to test against field data.

### 1.3. Complete spatial randomness case

Let us consider the similarity decay function (16). On the one hand, since the pair correlation function *g*(*r*) tends to 1 as *r* goes to infinity, χ(*r*) tends to a constant value asymptotically. On the other hand, if the complete spatial randomness hypothesis (CSR) holds for every species, then *g_α_*(*r*) ≡ 1 for every α, and the Sørensen index becomes again distance-independent and constant. In both cases we set

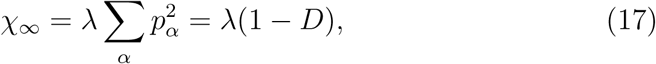

where 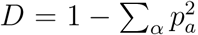 is the Gini-Simpson index [25, 26]. Hence χ∞ is the product of the probability λ of finding an individual times the probability 1 − *D* of finding two individuals belonging to the same species.

Formula (16) sets the range of applicability of our point process formulation of Sørensen index since it measures the average change of species composition at a scale which is comparable with the *largest of the* correlation range 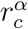 of the different species 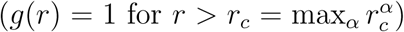 At a broader scale, the spatial point process description of the landscape composition becomes in a sense trivial, because all the species appear to be randomly distributed (CSR hypothesis).

Accordingly, the asymptotic value χ_∞_ does not depend on the clustering properties of the various species but only on their abundances. This feature may be useful for establishing a test of the theory independent of the chosen cluster model. Using the standard estimator for the intensity 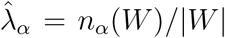, where 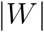 is the size of the whole study region *W* and *n_α_*(*W*) is the number of points of the α species within *W*, we derive the following estimator for χ_∞_ in (17)

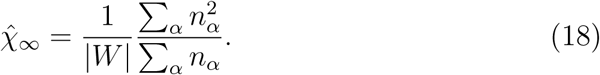

Let us suppose that the Species Abundance Distribution (SAD) φ(*n*) of the region *W* under study is known by other means, independently of the assumptions made for the spatial process. Then, equation (18) can be rewritten as

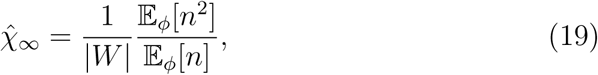

in which 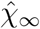 depends on the ratio between the second and first moment of *n*

### 1.4. Analytical formula for finite-size cells under CSR hypothesis

For later use, we derive here an analytical formula for the similarity between two regions of *finite and equal* area *a* under the CSR hypothesis, i.e. for a superposition of Poisson point processes (see e.g. [22]). Under this assumption, every point is uncorrelated to the others, hence the spatial point pattern is independent of the distance between points. This framework is thus intended to describe the similarity between two plots of finite area very far away so that species compositions are virtually independent. This is the approach taken in [18].

In this case, for two regions *A, B ⊂ W* of equal area *a*, setting *n_α_* = *n_α_*(*W*) we have –see (6), (7)–

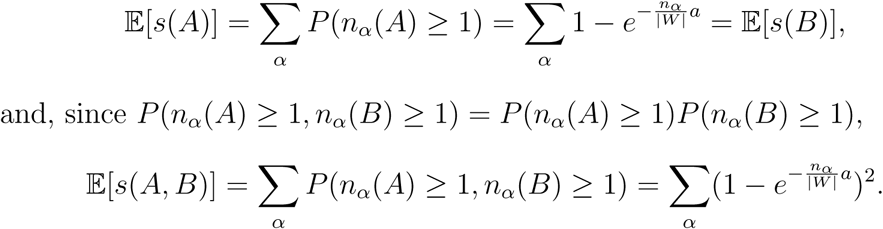

Therefore

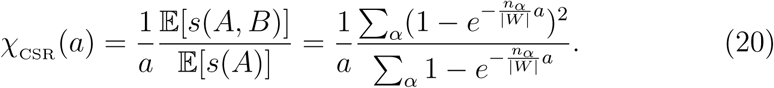

Formula (20) gives the asymptotic value of the similarity between two patches of finite area *a* far away (see pink solid line in Figure 3). This quantity can be considered as the discrete analogous of equation (10) in [18] under the CSR hypothesis (see also [19], Supporting Information F2).

Using the Taylor expansion of the exponential function *e*^*x*^ = 1 + *x* + *o*(*x*^2^), when *a →* 0 the above formula reduces to equation (18):

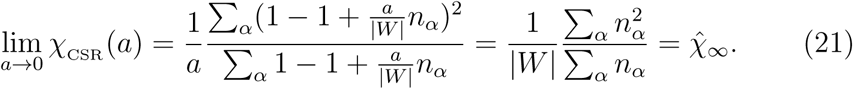

## 2. Estimators for χ and *g*

### 2.1. Direct estimators for χ

Let *S* be the total number of species and *N* the total number of individuals in our study region *W*, supposed to be a rectangular window of side lengths *l*_*x*_ and *l*_*y*_. For each species α, the coordinates 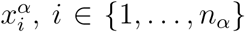 of all its *n*_*α*_ = *n*_*α*_(*W*) individuals falling within *W* are known.

In order to estimate the Sørensen similarity (8) whose average is (11), we first divide our region *W* in cells of area *a* and call 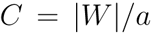 the total number of cells. Let *c*_*i*_ be the center coordinates of cell *i* = 1, …, *C*. Then, calling *K*_*r*_ the number of cells having distance *r* from each other, we give the following estimator

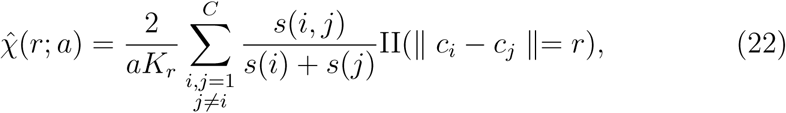

where *s*(*i*) is the number of species in cell *i*, *s*(*i, j*) is the number of species co-present in cells *i* and *j*, and II(·) is the indicator function:

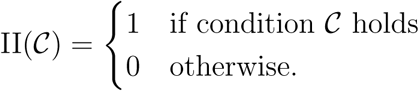

Let us remark that, with this estimator we are considering the *spatial* average over all cells in *W* having distance *r*, while point process theory considers the average, for fixed cells located in *x* and *y*, *over many realizations* of the point process. This latter cannot be computed, since, of course, we are given a single realization of the process, that is the ‘real’ forest. However, the two kind of averages (respectively over space and over realizations) are equivalent provided that the stationarity and isotropy hypotheses are realistic assumptions for our forest.

Denoting with *X* = *s*(*i, j*) and with *Y* = *s*(*i*) + *s*(*j*), we have that (22) will give an estimate of 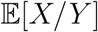. The estimator for the ratio 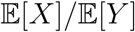 can, instead, be obtained through the following formula

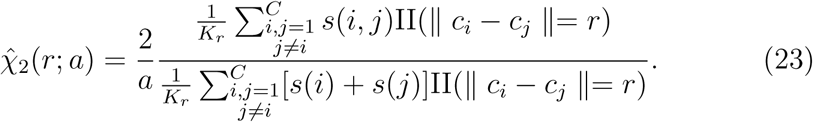

From (9), if we compute 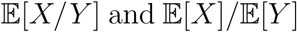 we can also compute *ϕ*(*x, y*). Let us note that, under the hypotheses of stationarity and isotropy under which we are working, *ϕ*(*x, y*) is a function of *r* = *x y*, i.e. *ϕ*(*x, y*) = *ϕ*(*r*). Using (22) and (23), we found that for our empirical data *ϕ*(*r*) *<* 0.05 at any distance *r* considered, leading to a relative error of less than 5% between these two estimators (see Figure 1) 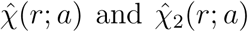 Therefore, in the following we will approximate *ϕ*(*r*) ≈ 0.

**Figure 1:**
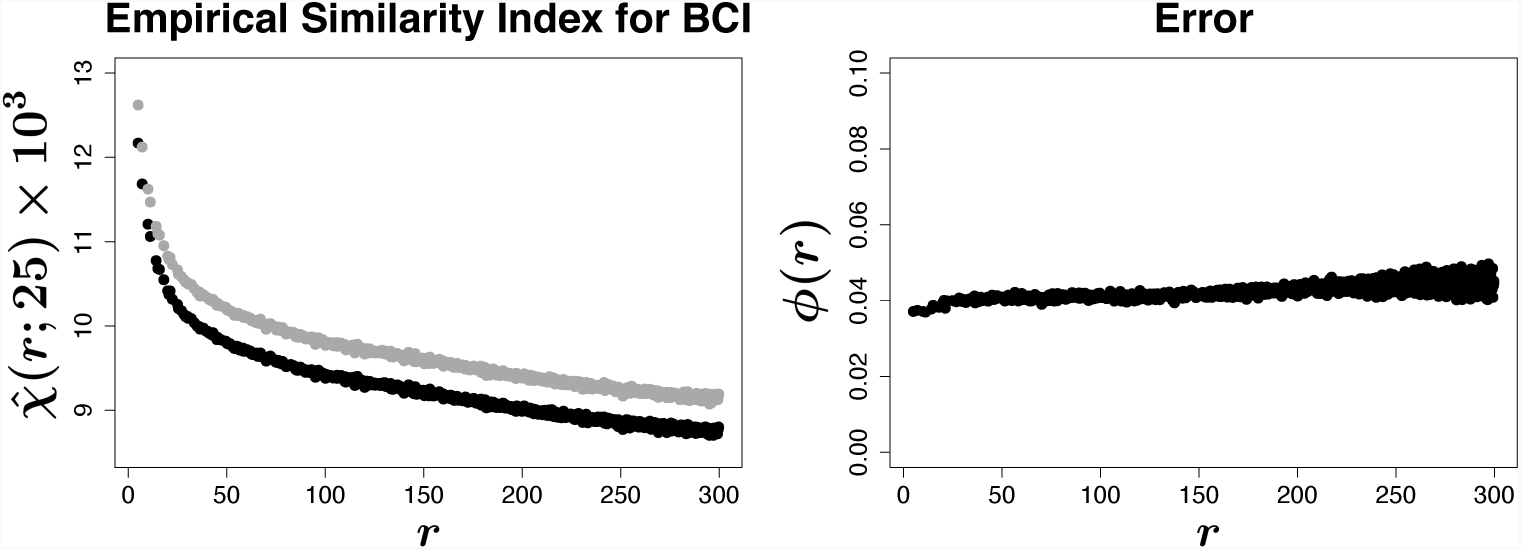
Comparison between empirical estimators of χ. We computed the similarity index for the BCI dataset considering species with more than 200 individuals through (23) (black curve on the left panel) and we compared it with the one obtained through (grey curve). In both cases we set *a* = 25. On the left we plotted the value of *ϕ*(*r*) at any distance. It resulted always smaller than 0.05, leading to a relative error between the two estimators of less than 5%. We therefore approximate *ϕ*(*r*) ≈ 0 in the rest of the paper.

### 2.2. Estimator for χ based on the estimator for g

Following [15] we estimate the empirical pair correlation function of the *α* species as follows

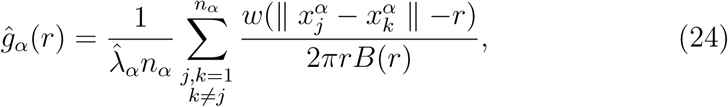

where 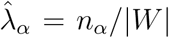 is the unbiased estimator of the density of individuals of the α species, || · || denotes Euclidean distance on the plane, *B*(*r*) = 1 − *r*(2*l*_*x*_+2*l*_y_ − *r*)/*l*_*x*_*l*_*y*π_ is the edge corrector function and *w* is the Epanenchnikov kernel, defined as

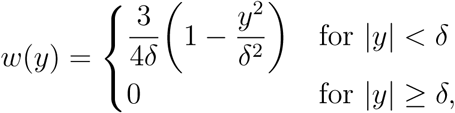

with 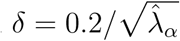

The estimator for the pair correlation function *g*_*X*_ of the superposed process *X* can thus be computed using formula (15) as

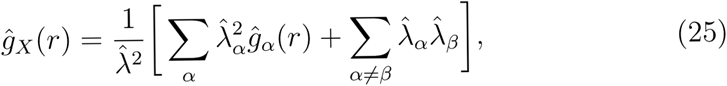

where 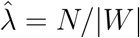 is the unbiased estimator of the total density of individuals. Therefore, an indirect estimator of χ can be obtained from plugging (25) above and (18) into (16):

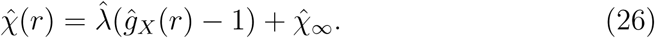

### 2.3. Finite cell size scaling

Note that the statistical estimator 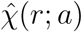 in (22) for χ(*r*) depends on an extra parameter, the cell size *a*. Since our analytical formula (16) is designed for an ideally infinitesimal area, we are faced with the problem of coupling the results of the direct estimator 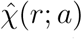 relative to finite area cells to the output of (26) where the finiteness is taken into account by the Epanenchnikov kernel. Below we show how to properly rescale the decay curves of 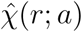.

In Sect. 1.4 we derived the analytical formula (20) for the similarity between two regions of equal area under the complete spatial randomness hypothesis. Out of this special case, it is hard to guess how the output of the similarity estimator (22) depends on the cell size *a*. We therefore tested the incidence of the cell size on the output of similarity estimator (22) by superimposing five different grids onto the 50ha plot, with square cells of area 1, 4, 25, 100 and 625 square meters respectively.

In Figure 2, we can see that the choice of the cell size *a* strongly influences the curves 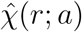 although the general trend results stable. However, as shown in the right panel of the same figure, the curves 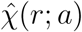 divided by their respective value at the largest considered distance, 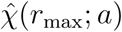, are *approximatively independent* on the cell size. For us *r*_max_ is the maximal available distance in the study area given by the shorter side of the rectangular study area, that is *r*_max_ = 500 m. In the following, we set 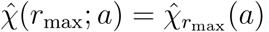.

**Figure 2:**
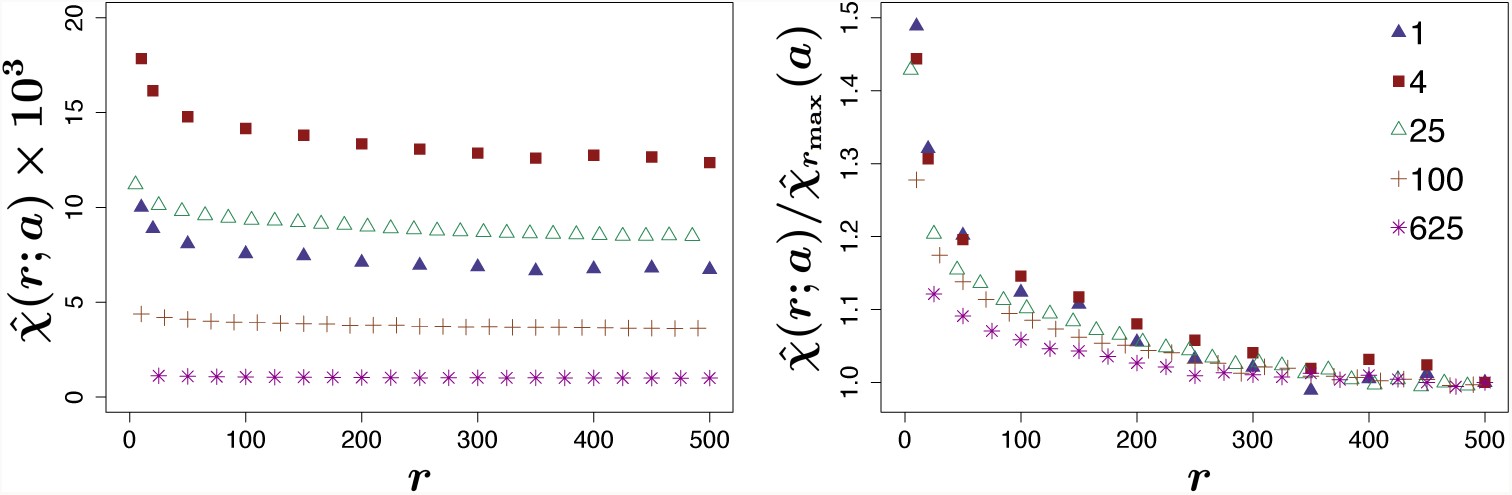
Sensitivity of Sørensen similarity index on cell size. We computed the similarity index for the BCI dataset considering species with more than 200 individuals. We superimposed to the 1000x500 observation window different regular square grids and estimated the corresponding Sørensen indexes 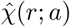 via (22). In figure, different colors represent different cell sizes, as in the legend. On the left panel: the choice of the cell size strongly influences the result, by “shifting the curve 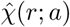 along the *y*-axis”. On the right panel: we divided each curve by its empirical value at the maximum considered distance, 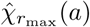. The resulting curve can be considered approximatively independent of the cell size and the error decreases with the distance.

Therefore, from Figure 2 right panel we can experimentally deduce that at *any distance r*

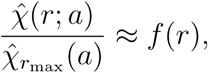

i.e. the ratio is independent of the cell size. Hence, for two cell sizes *a* and *b*

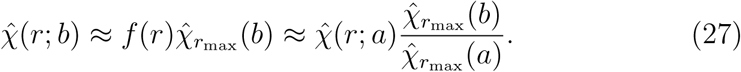

and taking the limit for *b* → 0 we may derive the statistical estimator for an infinitesimal cell size given the estimator for a finite cell size 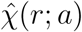

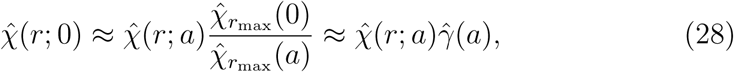

where we have set 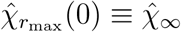 and

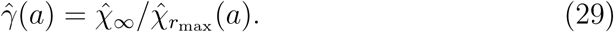

Note that the scaling factor 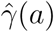 is the ratio of two similarities between plots very far away, where we may assume that the CSR hypothesis holds. In Sect. 1.4, we have derived the analytical function χ_CSR_ (*a*) of the similarity under CSR hypothesis and its limit for 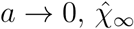 (see (20)).

We tested if the assumption 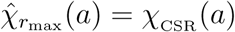 does hold for our study area so that we may compute *analytically* the scaling factor as

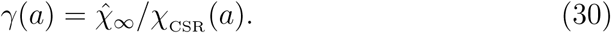

Results are contained in Figure 3, where we plotted the function χ_CSR_ (*a*) defined by (20), for *a* up to 625 square meters (solid pink line) and the empirical values of 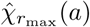 computed through the estimator (22) for different cell-sizes (colored symbols). There is a very good agreement between these two quantities, which improves with the cell size *a* (the off-curve point *a* = 1 square meters is probably due to the finite diameter of the plants) meaning that, at least for *a ≥* 25 square meters, the CSR hypothesis holds for distances *r ∼ r*_max_ = 500 meters where correlations between points become negligible. Note that the curve χ_CSR_ (*a*) (pink solid line) tends asymptotically to 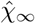 (blue line) for *a* → 0 as prescribed by (21).

**Figure 3:**
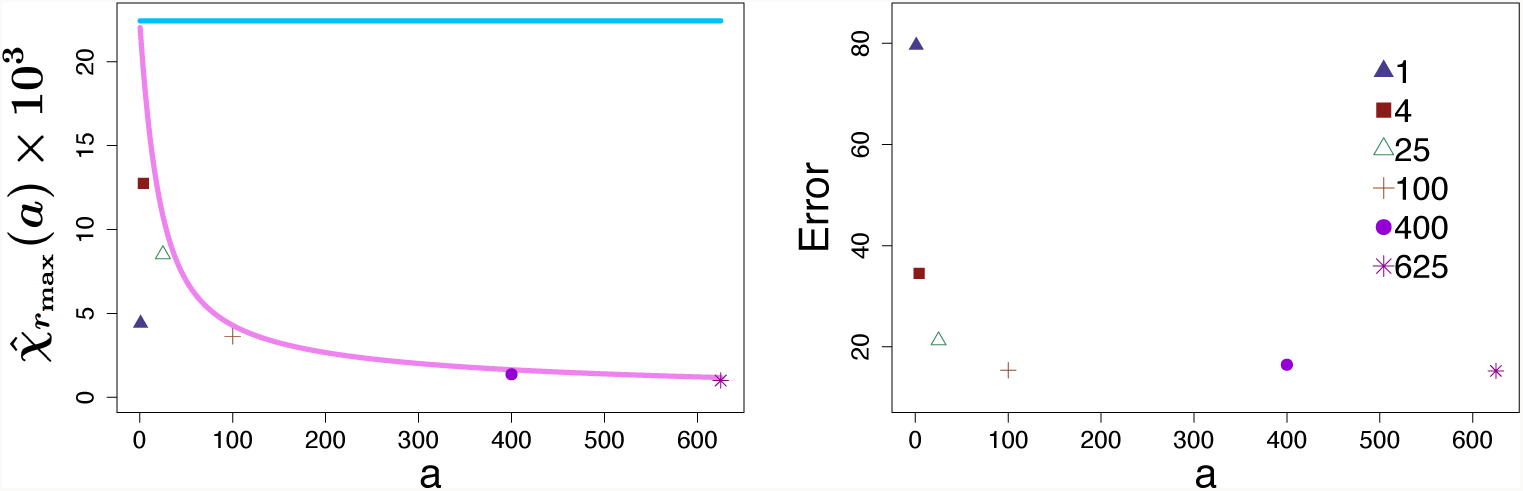
χ_∞_ for finite-size cells. On the left: the pink line represents the asymptotic value of the theoretical similarity index, χ_CSR_ (*a*) as a function of the cell-area – see (20)–. Colored symbols are the empirical values 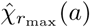 computed via (22) for cell sizes of 1, 4, 25, 100, 400 and 625 square meters, respectively, and considering only species with more than 200 individuals. The straight light blue line represents the value of 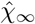 for infinitesimal cells computed through the abundances (see (18)). On the right: relative percentage error between the empirical values and the theoretical ones.

With the scaling formula (28), we can now rescale the output of the direct statistical estimator 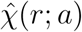 for finite cell size *a* to an infinitesimal cell size in order to compare it with the output of the indirect estimator (26) based on the pair correlation function *g_X_* of the superposed process.

Note that the rescaling (both upscaling or downscaling) can be done also between two empirical curves 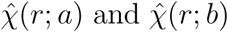 through formula (27), using the empirical scaling factor 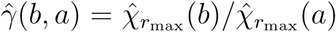 or the analytical one γ(*b, a*) = χ_CSR_ (*b*)*/χ*_CSR_ (*a*).

In Figure 4 we downscaled the similarity decay function 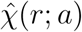 estimated via (22) with *a* = 625 to 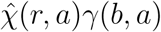 for *b* = 25, 100, 400 (dashed lines) and compared the curves with the original ones 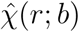 for the same values of *b* (colored symbols). As expected, for such fairly large areas, all the rescaled estimators are in good agreement. In the sequel we used the empirical scaling factor 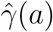 for smaller plots (1 to 4 square meters).

**Figure 4:**
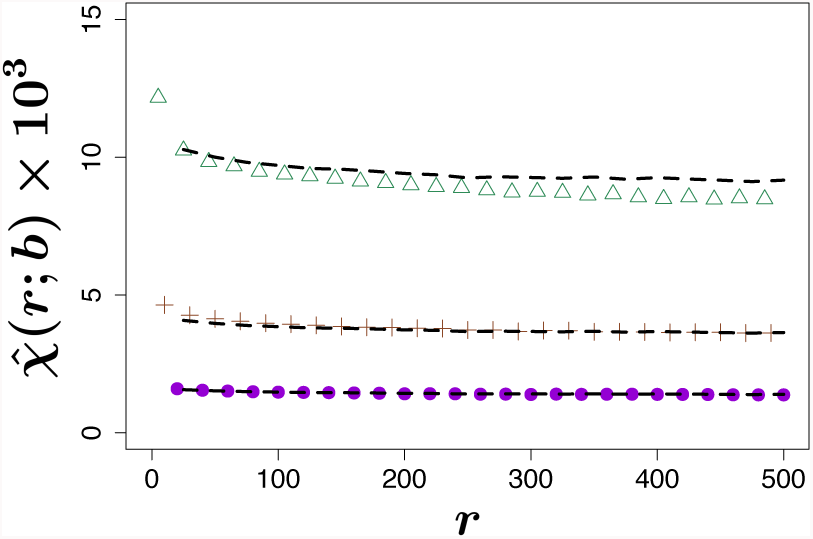
Comparison between rescaled χestimators. We tested the goodness of rescaling (27) by plotting the estimator (22) for 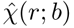 with *b* = 25 (triangles), *b* = 100 (crosses) and *b* = 400 (dots) against the rescaled one 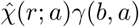 for *a* = 625 (dashed line). The difference between the two curves increases with |*b − a*|, but there is always an excellent fit.

## 3. Study of clustering using Neyman-Scott processes

By construction, the Sørensen similarity decay function (16) depends on the relative abundances of species and on their pair correlation function. This latter depends essentially on the clustering of the individuals. Therefore, the crucial point for the description of real data patterns is the choice of the cluster model of the point process. The theory exposed in this section, establishing a link between the form of the dispersal kernel -see iii) below– and the form of the resulting similarity decay function, could help to select the form of the dispersal kernel from the empirical similarity curve as an inverse problem. In particular, it would be interesting to determine the dispersal kernel that gives rise to a compound exponential decay function, as predicted by Hubbell neutral theory. The reader interested to the application of the theory exposed above to real forests can go directly to Sect. 4 below. In this paper we limit ourselves to Neyman-Scott (NS) processes [28].

These processes have found large applications in ecological theory due to their ability to model the clumping mechanism of plants’ species in which daughter seeds are spread around a parent tree’s location. They are the result of a three steps procedure:

i. Parent points are distributed according to a homogeneous Poisson process with intensity ρ.
ii. To each parent, a random number of offspring is assigned, drawn from a Poisson distribution of intensity *µ*.
iii. The offspring are identically and independently scattered around their parents with a fixed spatial probability density given by a *radial* function *dγ*(*r*), the so-called *dispersal kernel*, depending on some parameters γ.

The resulting process is formed only by the offspring’s locations. Independently of the offsprings’ distribution, the intensity function of the process is given by the product λ = ρµ. The pair correlation function of a NS cluster process is given by (see [29], pp. 318-319)

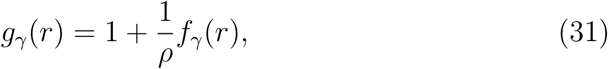

where *f*_γ_(*r*) is the convolution of the 2-dimensional probability density *dγ*(*r*):

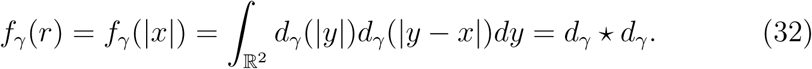

Note that *f*_*γ*_(*x*) is the probability that two offsprings belonging to the same cluster have *x* as the vector difference of their positions with respect to the cluster center. If *d*_*γ*_(·) is a radial function, so is *f*_*γ*_(·).

Considering *S* Poisson cluster processes *X*_*α*_ of parameters (*ρ*_*α*_, *µ*_*α*_, *γ*_*α*_), the Sørensen index of similarity of the process resulting from their superposition can then be obtained by plugging formula (31) into (14)

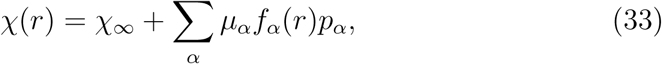

where we have set *f*_*α*_ = *f*_*γα*_ for simplicity. In the general case, the form of the function *d*_*α*_(*r*) reflects the cluster characteristics and may have short or long tails. The most used are Gaussian (single or mixture), inverse power or exponential (see [30]). Since *χ*(*r*) tends to χ_∞_, necessarily

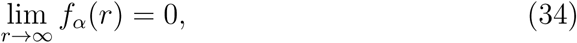

while the limit of *f_α_*(*r*) for *r* → 0^+^ may be finite or infinite (in this latter case the function is said to have a pole at 0). Our aim now is to compare the above curve (33) for χ(*r*) with the empirical ones (22) and (26) coming from field data to select the best cluster model and determine the corresponding cluster parameters.

### 3.1. Dispersal kernels functions

In the Supplementary Material we report in details the theory necessary for computing the convolution function *f_γ_* given the dispersal kernel *d_γ_*. Here we give the results.

a) *Exponential cluster*. For a 2-dimensional radial probability density of the form

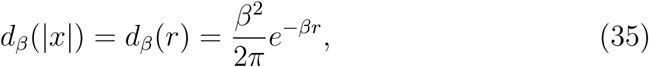

the computation gives

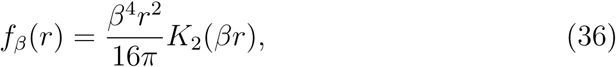

where *K*_*n*_(*z*) is the modified Bessel function of second kind [31]. Note that, for *z →* 0, it holds that 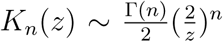 Hence *f* (0) is a finite quantity.

The average of *r* with respect to *d*_*β*_(*r*) gives the average cluster radius that is the average distance from parent point. For an exponential cluster this is 2*/β*.

b) *Gaussian cluster*. Another famous example of Poisson cluster process is the modified Thomas point process [32, 19, 33, 34, 35], where the cluster probability density is given by a bivariate Gaussian

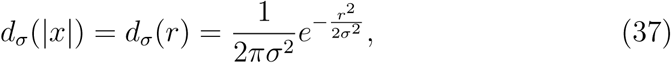

whose convolution is

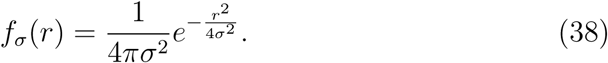

The average cluster radius in this case is 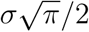.

c) *Gaussian mixture* (also called bivariate Student *t*, 2*Dt* in the following).

These are obtained as a continuous mixture of Gaussian kernels with variance parameter distributed as the inverse of a Gamma distribution (see [36, 20]). The 2-dimensional radial probability density depends on two parameters *b* > 0 and *p* > 0

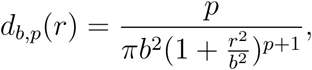

giving the convolution function

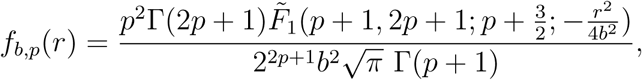

where 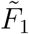 is the hypergeometric 2*F* 1 regularized function. Note that the average cluster radius is

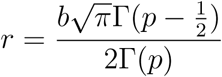

showing that *b* is a scale parameter and that the average radius is *infinite* for *p* ≤ 1/2. Indeed the radial density *d*_*b,p*_(*r*) has a fat tail (a power-law) hence this type of cluster is well suited for describing species with long range intra–specific correlations.

d) *Cauchy cluster*. The special case 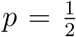 (2-dimensional Cauchy density) has a 2-dimensional radial probability density of the form 2 depending solely on *b*, and has a convolution function of the form

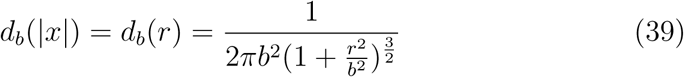

depending solely on *b*, and has a convolution function of the form

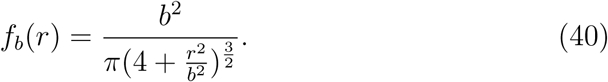

The choice of the dispersal kernel *d*(*r*) is a crucial point also in seed dispersal studies. For an overview of the problem see e.g. [36, 20]. The validation of the dispersal kernel choice is particularly difficult for tropical rainforests where the tree crowns overlap and therefore it is impossible to identify the seed shadow dispersed by a single tree.

### 3.2. Preliminary test on computer-generated forests

We tested the validity of estimator (22) rescaled according to (28) on four artificial forests generated according to different point processes: a Poisson one and three cluster processes (exponential, Gaussian and Cauchy). In all cases, we considered a square window of side 500 meters and generated a forest consisting of 50 species with abundances distributed according to a normal distribution of mean and standard deviation equal to 1000 and 300 individuals, respectively.

For the exponential and modified Thomas cluster processes, to each species was assigned a random average radius *r* drawn from a normal distribution of mean 20 and standard deviation 5. These values determined the cluster parameters 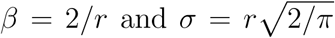. Since the average radius is not well defined for a Cauchy cluster process, in this case we arbitrarily set the cluster parameter *b* equal to *r/*2.

Once generated the four forests, we estimated the similarity decay function for infinitesimal area using the rescaled estimator 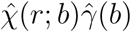 for *b* 25, 100, 200 and 625. We then compared the empirical curves with the theoretical similarity decay functions given by (17) for the Poisson process and (33) for the Neyman-Scott processes. The agreement between the empirical and theoretical curves increases as cell size *b* decreases, but it is always very good (see Figure 5).

**Figure 5:**
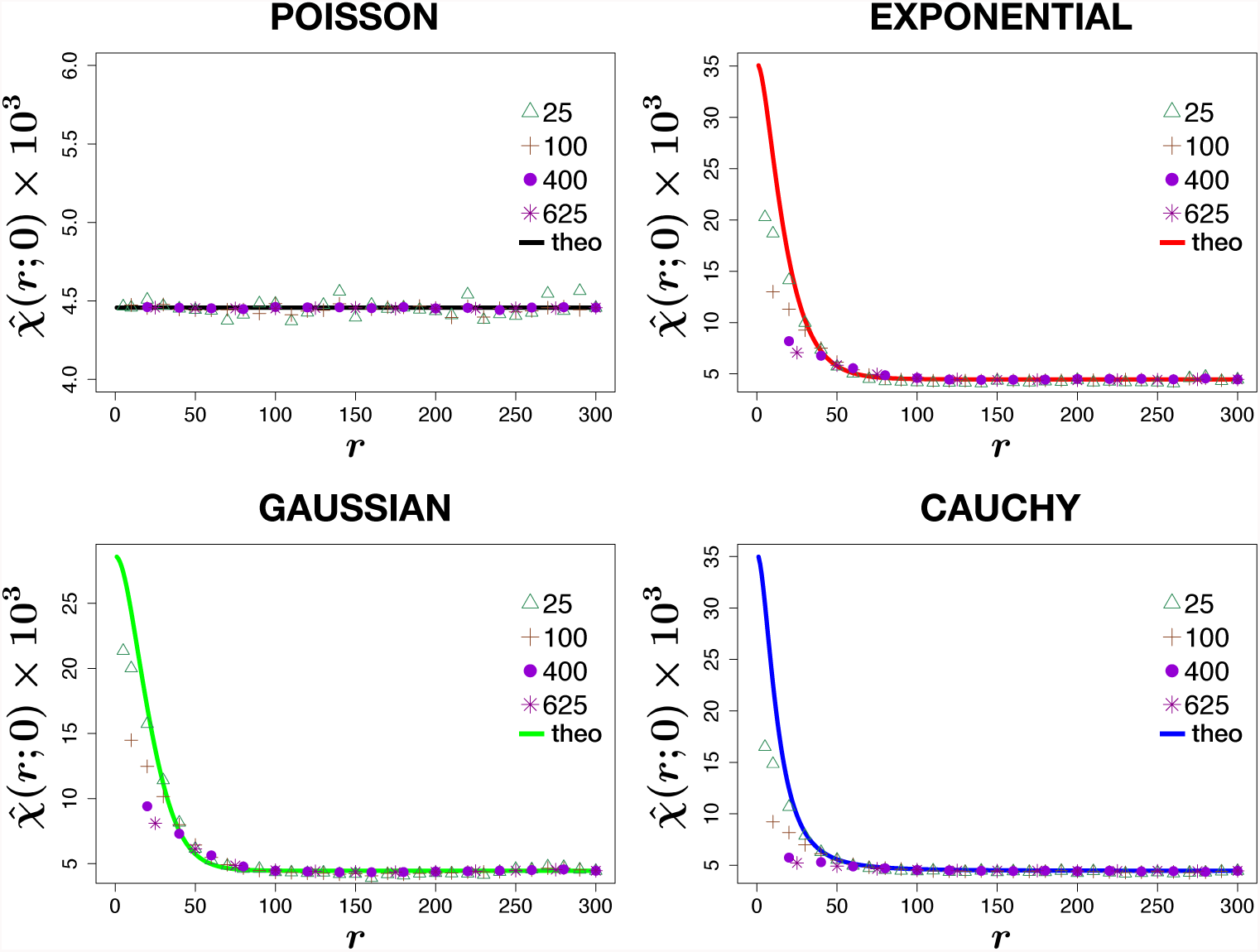
Test of estimator (28) on four artificial forests generated according to different point processes: a Poisson one and three cluster processes (exponential, Gaussian and Cauchy). We compared 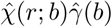 for *b* = 25 (triangles), 100 (crosses), 400 (dots) and 625 (stars) with the theoretical similarity decay function (solid curves) given by (17) for the Poisson process and by (33) for the Neyman-Scott processes. The agreement between the empirical and theoretical curves increases as *b* decreases, but it is always very good.

## 4. Test of estimators on BCI ecological dataset

We test our analytical formula and its related estimators on the Barro Colorado Island ecological dataset (BCI) consisting of the spatial coordinates of 222602 individuals belonging to 301 different species of plants within a 50ha rainforest plot.

### 4.1. Species selection and subsampling

Species selection. To see if neglecting the scarcely abundant species strongly affects the similarity index for BCI, we superimposed a 5x5 grid onto the plot and compared the distance-dependent Sørensen index computed via equation (22) taking into account species having least abundance of 0, 20, 100, 200, 300 and 500 individuals, which represent, respectively, the 100%, 73%, 49%, 36%, 30% and 24% of the total biodiversity and which account for the 100%, 99%, 98%, 96%, 93% and 90% of the total number of individuals. From our analytical formula, we know that the similarity is affected only by the most abundant species.

On the left panel of Figure 6 we plot the obtained empirical curves, while in the right plot the corresponding relative percentage error with respect to the Sørensen index computed for the whole forest. By selecting the species with a population of at least 200 individuals (107 species), we get a curve which differs form the similarity curve of the whole BCI for about 5%, a reasonable restriction for our goal.

**Figure 6:**
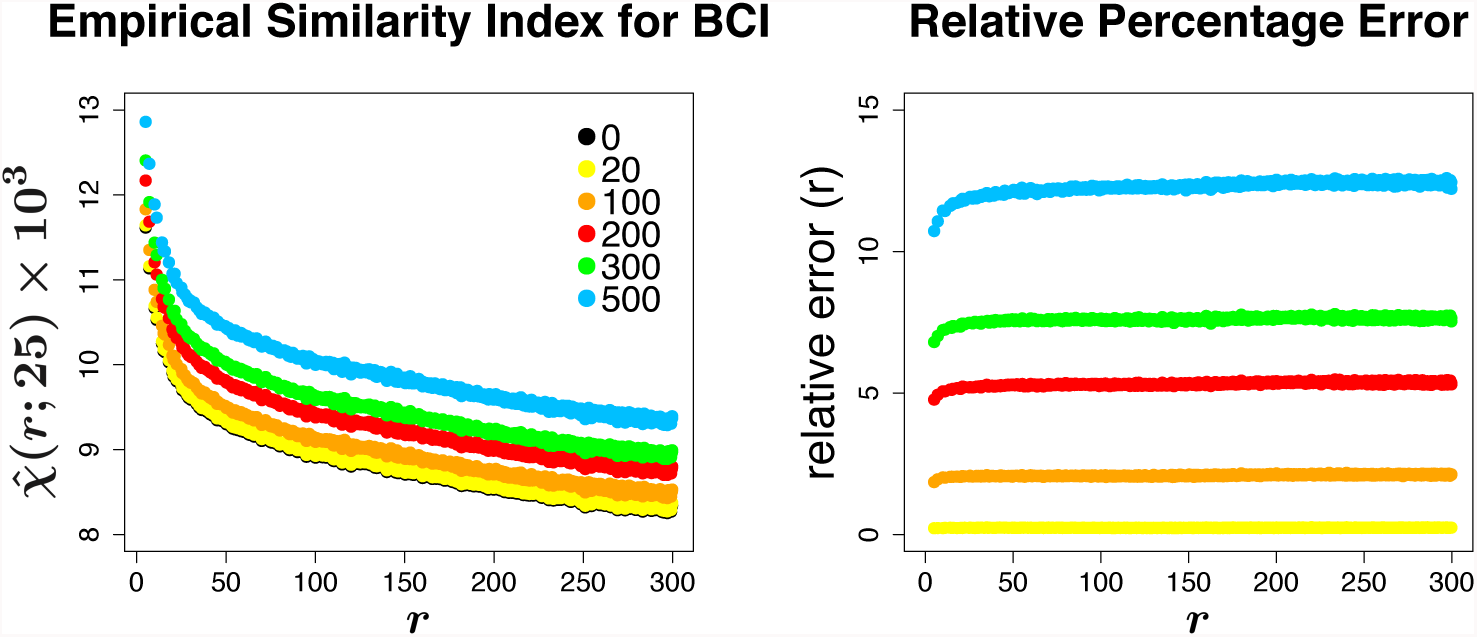
Sensitivity of Sørensen similarity index on abundances. We have already noticed that our distance-dependent Sørensen index is dominated by the most abundant species. On the left panel, we show the different curves 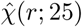 that we obtained via estimator (22) by taking into account only species having least abundance as indicated in the legend. On the right panel, we computed the relative percentage error with respect to the black curve, which is the similarity index for the whole forest.

Sub-sampling. As a last preliminary test, we checked whether sub-sampling affects the estimate of the Sørensen index. Again, we superimposed a 5x5 grid on our study region and considered three scales of sub-sampling by ran domly taking the following percentages of the 20·000 available cells: 50%, 25% and 5%. In Figure 7, points are the result of averaging over 10 trials for each sub-scale. We can observe that, although lower percentages affect the curve by significantly increasing the fluctuations, the general trend of the curve is very well preserved for a sub-sampling of up to 25% of the data.

**Figure 7:**
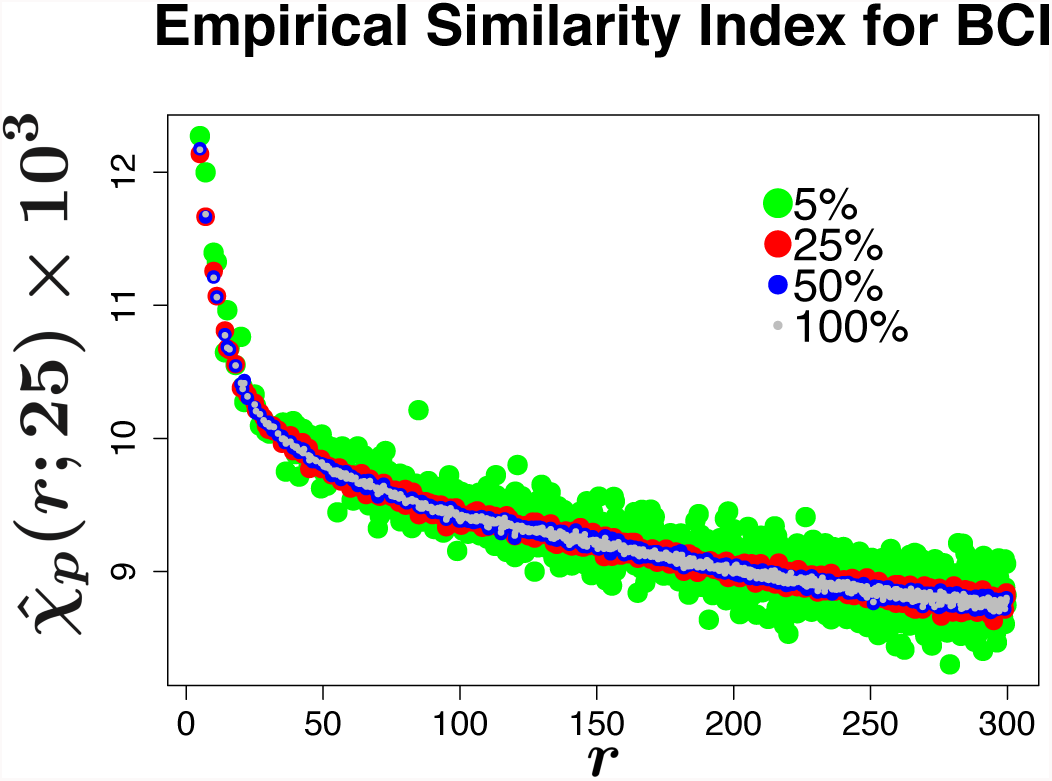
Sensitivity of Sørensen similarity index on sub-sampling. We investigated the effect of sub-sampling in computing the empirical Sørensen index estimated through (22). We first superimposed a 5x5 grid on the window, corresponding to *C* = 20·000 square cells. For every percentage *p* shown in the legend, we randomly choose *pC* cells and compute the distance-dependent similarity index between them, 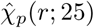 Points are the mean on 10 trials. Clearly, the lower the percentage of considered cells, the more fluctuations affect the curve, although the trend results very stable under subsampling.

### 4.2. Comparison of direct and indirect similarity estimators

We can test the validity of our similarity decay model (formulae (14) and (16))

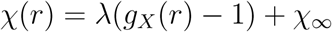

by estimating independently its left and right sides. For χ(*r*) we use the estimator 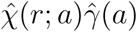 given by (22) multiplied by the finite cell size scaling factor 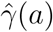 in (29), whereas for the right hand side we use the indirect estimator (26). Results are displayed in Figure 8. We can see that there is a very good agreement between the two estimates if the smallest cell sizes are used (*a* = 1 or 4 square meters), which are the closest to the theoretical hypothesis of infinitesimal cell size, but the use of the finite cell size scaling factor gives fairly good results even for greater cell sizes. Moreover, we see that the estimated decay curves tends to the analytical one monotonically

**Figure 8:**
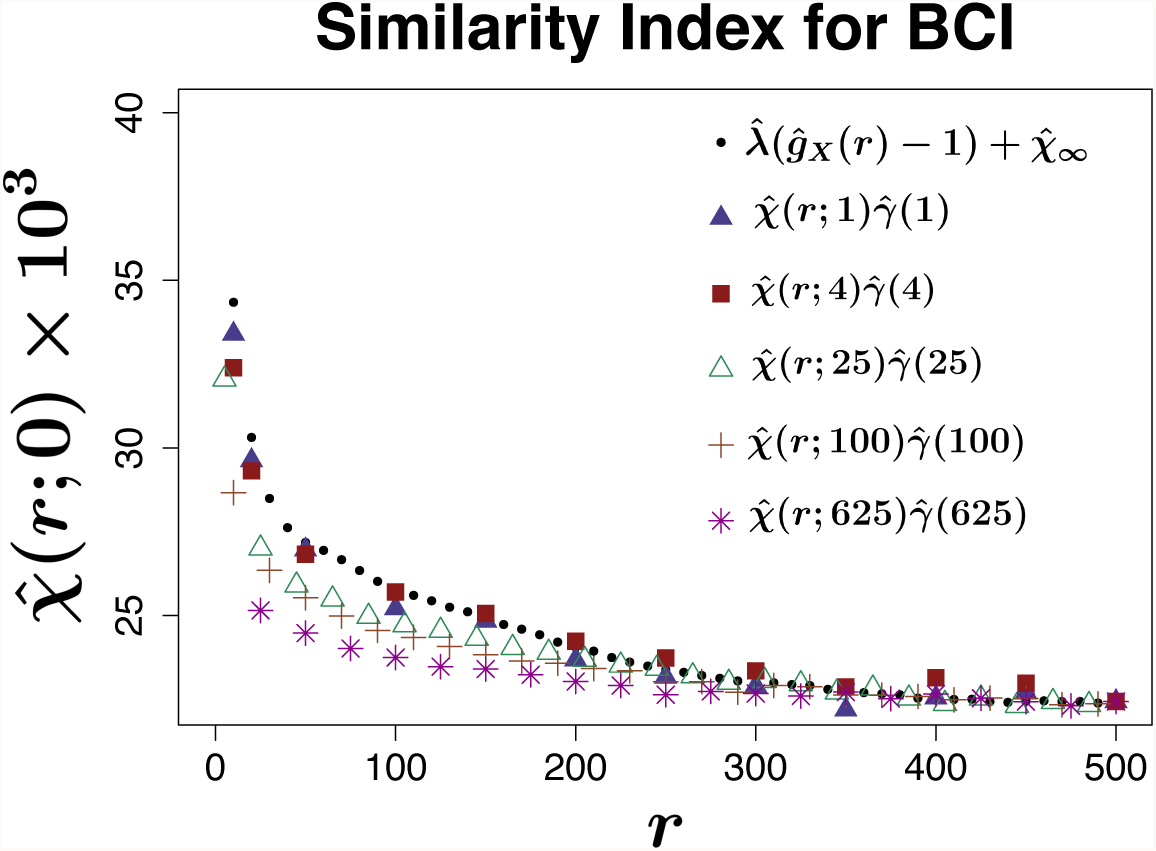
Similarity Index for BCI. Similarity index for BCI dataset computed by (26) (black dots) compared with the estimator (28) for different cell sizes *a* (colored symbols). The agreement between the two estimators increases as the cell size decreases. This is in accordance with the theory.

from below. Therefore the analytical curve sets an upper bound for the density of similarity. Note also that when using the smallest cell sizes the curve displays a tri-phasic behavior with a steep initial descent, a linear descent in the middle and a hollow tail. This behavior is not captured with coarser cell sizes. We will discuss a possible explication of this phenomenon in Sect. 4.3. We stress that the basic estimator (22) is independent of any assumptions on the clustering of the individuals, which are not known a priori, while these assumptions are contained in the indirect estimator (26). The good agreement between the two estimates supports the conclusion that, at the considered scale 0 *< r <* 500 m, clustering is a main driver of species turnover. Also, at contrast with Hubbell Neutral theory [11], we find that rare species do not contribute significantly to the specie turnover even at local scales.

### 4.3. Comparing estimated and theoretical similarity functions

Let us now focus on modelling BCI species through the three NS point processes described in Sect. 3.1: exponential, Gaussian (modified Thomas) and 2*Dt* kernel processes.

For each species α, the first step is to estimate the set of parameters (*ρ*_*α*_, *µ*_*α*_, *γ*_*α*_) which best describe its pattern. We do this by the method of minimum contrast [32], which relies on the minimization of the following integral

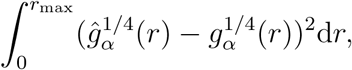

where 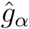 is the empirical pair correlation function estimated according to (24) and *r*_max_ is the maximum considered distance, equal to 500 meters.

By inserting the fitted parameters of each model into the corresponding formulation of equation (33), we can get the predicted theoretical similarity curve for the BCI forest (see Figure 9). We find a good agreement between the model prediction and the empirical data for all cluster types. Nevertheless, the Cauchy cluster type gives the best results at fitting the single species pair correlation function (see the results of the χ^2^ test in the Supplementary Material).

**Figure 9:**
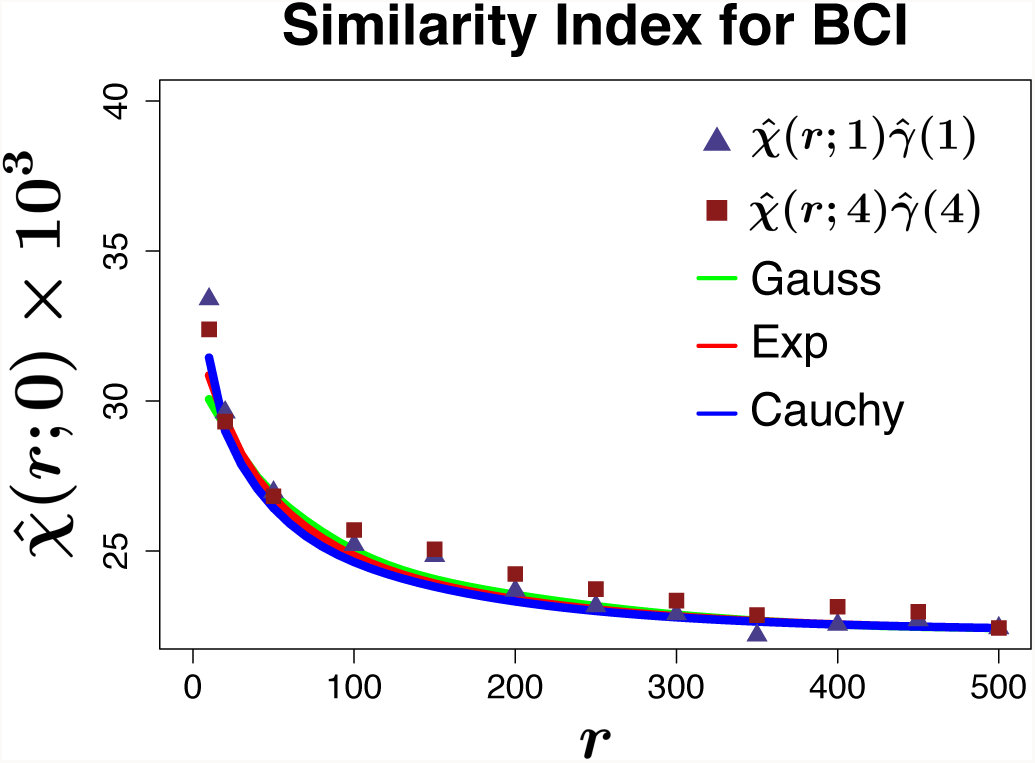
Sørensen index for BCI. Comparison between the empirical distancedependent Sørensen index 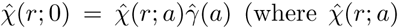 has been computed via (22) and 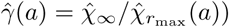 for square cells of area 1 (triangles) and 4 (squares) square meters and the exact functional form of χ(*r*) by the three cluster models using (33). We find a good agreement between model prediction and empirical data for all cluster type.

Our analytical similarity decay function (16) was derived using the point process framework and is based on the following assumptions: 1) we compare the species’ composition of two regions of *infinitesimal size* and 2) the point process is *stationary and isotropic* (i.e. translation and rotation invariant) and *homogeneous*, i.e. the *intensity λ*, which is the density of individuals, is constant (see Sect. 1.1). Here we would like to discuss the impact of hypothesis 2).

To investigate this, we generated three artificial forests as follows: for each BCI species having more than 200 individuals, we generated a Neyman-Scott *homogeneous, stationary and isotropic* cluster process within the 50ha plot having the same number of individuals as the original species and according to the three different cluster types parameters. We then computed the empirical Sørensen similarity index 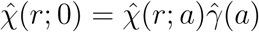, with *a* = 2 and 25 for the superposed process and compared it with the theoretical one (equation (33)). Results are displayed in Figure 10.

**Figure 10:**
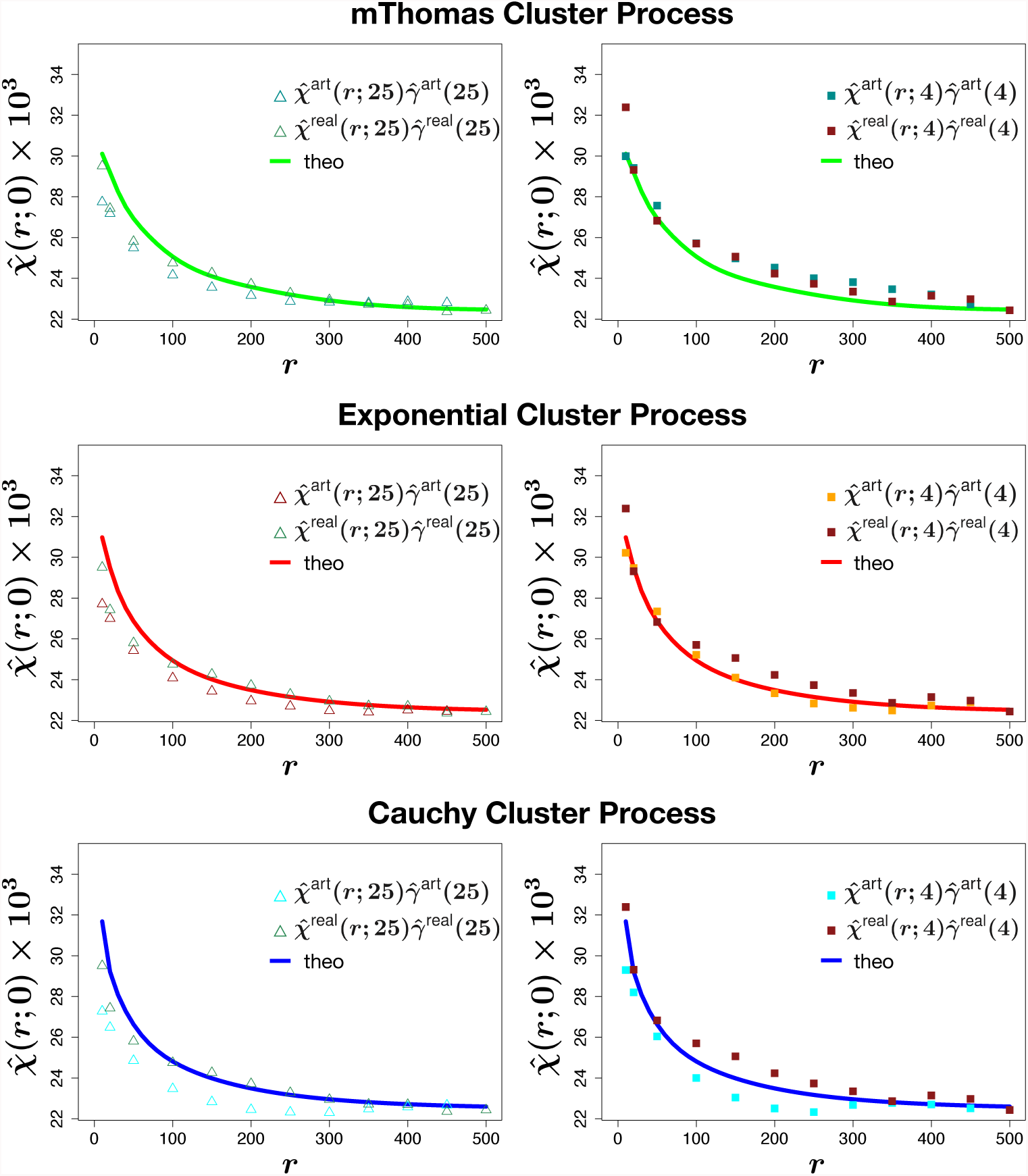
Empirical Sørensen index for artificial forests. We generated three artificial forests as follows: for each species of BCI having more than 200 individuals, we generated a Poisson cluster process (modified Thomas in the top panels, Exponential in the middle panels and Cauchy in the bottom panels) within the 50 ha plot having the same number of individuals as the original species. We then computed the empirical Similarity index for the new generated superposed process 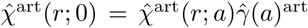 (the superscript “art” stands for artificial forest) and compared it with the theoretical one (see (33)) and the empirical one for the real BCI 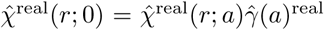 (the superscript “real” stands for real forest), when using 25 square meters cell area (left column) and 4 square meters cell area (right column).

For the forests generated according to the three cluster model hypothesis 2) holds. Considering a cell area of 25m^2^ (left column), the empirical similarity decay function of the artificial forest does not display the linear descent in the middle part, which is on the contrary present in the empirical curve of the BCI data. At smaller cell area scale(4m^2^, right column), we found mixed results: the linear descent is present in the Gaussian dispersal kernel, is absent in the exponential one and is very hollow in the Cauchy kernel. This is a consequence of the different average cluster radius of the three models, which is infinite for the Cauchy cluster.

To give a hint for a possible explanation for the linear descent phenomenon in Figure 11 we show the pattern of two different species of the BCI forest, one showing an homogeneous behavior (left column, first panel) and one affected by anisotropy and non-stationarity (right column, first panel). For both species, bottom panels show the analytical curves of the pair correlation function (colored lines, see (31)), with parameters obtained by the minimum contrast method against the empirical pair correlation estimated via (25). In contrast to the first species, where all three models are able to capture the empirical curve, for the second species the fit is much worse and, more importantly, the curve shows a hollow part in the middle due to overdispersion at that scale. Since the pair correlation function of the superposed process is the sum weighted by the abundances of the pair correlation function of the different species, we may think that the linear descent of the curve in its middle part is an average behavior due to the fact that for some species the patterns are not homogeneous nor translation or rotation invariant.

**Figure 11:**
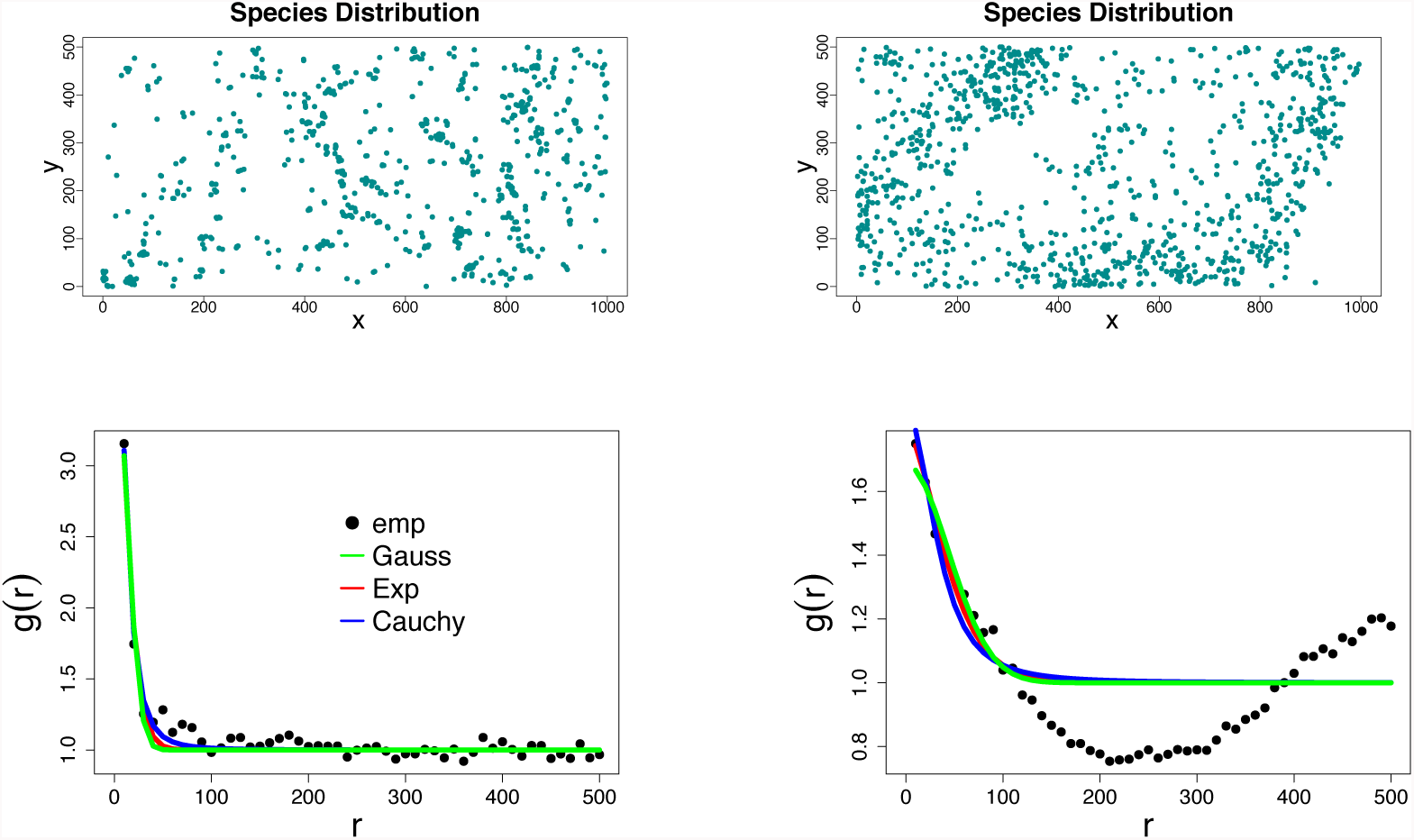
Pair Correlation Function for two species. On the top: two species distributions within the 1000x500 surveyed area of the BCI. We can notice that the species on the right panel shows a non-isotropic nor homogeneous pattern, resulting in contrast with our model hypothesis. Such behavior does not characterize the species on the left. On the bottom: empirical pair correlation function computed via (24) (black dots) and the analytical one (solid lines) computed through equation (31) with parameters fitted by minimum contrast method.

## 5. A synopsis of similarity decay functions

In this section we give an account of the various approaches to the problem of describing the decay in similarity with the distance that we have found in literature. In their pioneering paper [9] Nekola and White studied North America boreal spruce forests using data from 34 nine hectare plots distributed from Newfoundland to Alaska. The similarity was computed using Jaccard index and species were subdivided in homogeneous classes in terms of growth or dispersal form. Linear regression was used to calculate the decay rate of the logarithm of the similarity against linear distance. This implies an *exponential* rate of the distance decay, with different exponents for various classes.

In Hubbell’s neutral theory of ecology [11] the similarity decay is also considered for an artificial community. The form of the decay function is a *compound exponential*, i.e. a linear combination of exponentials with different exponents. The steeper decay rate is the contribution due to the rare species, which are also confined to restricted areas, while the tail of the curve has a lower decay due to the abundant and widespread species with lower turnover. The overall decay is steeper and the overall similarity is lower if a smaller grain size (i.e. plot size) is used. This latter aspect can be discussed only qualitatively within the theory. However, the dependence of the decay curve on the size of the plot is an unavoidable consequence of the very definition of similarity, which is area-dependent. This renders more difficult the comparison of different graphs realized with diverse grain sizes and extents, and its potential impact on the conclusions drawn from these data have been recalled in various works [24, 27].

In [19], an analytical model for the similarity decay function is presented, extending a spatially implicit model contained in [18]. This spatially explicit model considers two small regions *A* and *B* of area *a at distance r* and is based on the conditional probability

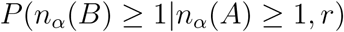

that gives a distance dependent similarity index. In [19], the probabilities are computed for a specific spatial point process, a modified Thomas cluster process of parameters (*µ, ρ, σ*). The resulting formula of the Sørensen similarity, for a discrete number of species (see [19], Supporting Information F2) is the following

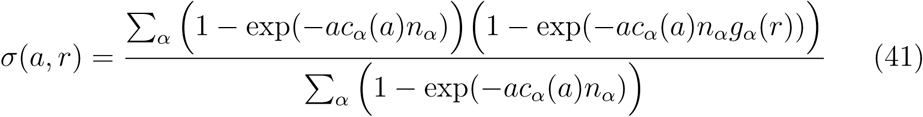

where *g*_*α*_(*r*) is the pair correlation function of the modified Thomas process

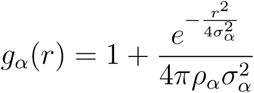

and *c*_*α*_(*a*) is an area-dependent correcting factor for the clustering of individuals having dimension *area−*^1^:

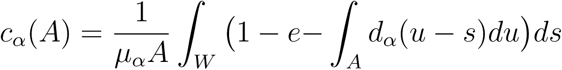

where *d*_*α*_ is the Gaussian dispersal kernel function (see (37)). For randomly distributed individuals *c*(*a*) = 1, while *c*(*a*) tends to zero for highly clustered patterns. Note that, for a random pattern, also *g_α_*(*r*) = 1 for every specie α. Therefore, in this case, dividing (41) by the area *a*, we obtain (20). For the continuous case the formula is more involved. However, when the term *ac_α_*(*a*)*n_α_* is small, keeping only the leading term in the Taylor expansion of the exponential as done before, we find that the Sørensen similarity for very small plots is (this derivation is ours)

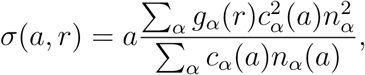

which again, under the random placement hypothesis, reduces to (21) when divided by *a*. There are probably other approaches to the problem of determining the form of the similarity decay function of which we are not aware of, but, as far as we know, it seems to us that formulating the theory for the similarity between plots of finite area produces very complex formulae in which the dependence on the area is not easy to investigate. We are convinced that the formulation presented in this paper based on infinitesimal plots offers a clearer picture of the problem.

## 6. Conclusion

Spatial point processes are a powerful statistical tool for the description of patterns in tropical rainforests. In particular, we investigated the role of spatial clustering in shaping the curve of species turnover with the distance. Therefore our results can be applied to a forest extent where climatic, orographic or other shaping factors are not present. In this framework we have derived an analytical formula for the average decay in similarity with the distance between two relatively small plots. A peculiar trait of our approach is the use of the *spatial density* of the similarity with respect to the area. Essentially, we find that the decay function of the similarity density is given by the pair correlation function of the whole forest (see (16)) and that it is determined by the most abundant species. This formula thus establishes a link between a very important concept in quantitative ecology like the decay in similarity, with a widely used concept in the statistical description of a general particle systems. This hollow curve tends to an asymptotic value which is determined by the relative abundances. Our similarity decay function χ(*r*) is related to the distance-dependent Simpson index of Shimatani [1], β(*r*), and to the codominance index *F* (*r*) in [20]. To test the analytical theory against real data, we have designed a statistical estimator for the similarity which is based on presence-absence counts on plots of *finite* size and on an area-scaling factor. We are thus able to interface our analytical theory, which refers to plots of infinitesimal size, with the estimator.

The limiting hypothesis of relatively small size of the plots with respect to the distance between them is present in all other works we examined dealing with this problem, and it is not easy to manage (see e.g. [24]). We think that in our approach the dependence on the area is easier to control since it is transferred to the statistical estimator, which is flexible enough. We tested our findings on the extent of the study area of BCI and Pasoh forests, which are both of 50ha, obtaining a very good agreement with the empirical data. At larger scales, if other drivers of biodiversity other than clustering of individuals are acting, our model can not be applied directly. Nevertheless, if no strong environmental inhomogeneities are encountered, it could be effectively employed for larger portions of rainforests, where a complete survey of all individuals is impossible, since it performs well with respect to random subsampling.

## Acknowledgments

We thank the Center of Tropical Research Science (R. Condit, S. Hubbell, Foster) for providing the empirical data of the BCI and Pasoh forests. We thank the anonymous referees for their valuable comments and suggestion on the presentation of the contents of this work.

## Supplementary Material

The distance decay of similarity in tropical rainforests. A spatial point processes analytical formulation

### 1. Convolution of 2-dimensional radial functions

We briefly report here the theory necessary for computing the convolution function *f* given the dispersal kernel *d* [1]. Given a function *f* (*x*) on the plane, 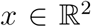, let us denote with 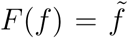 its Fourier transform. We recall also the Convolution Theorem:

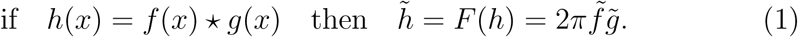

For radial functions *f* (*r*) = *f* (|*x|*) the Fourier transform can be computed^1^ using the Bessel function *J*_0_(*z*)

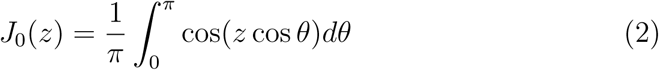

as

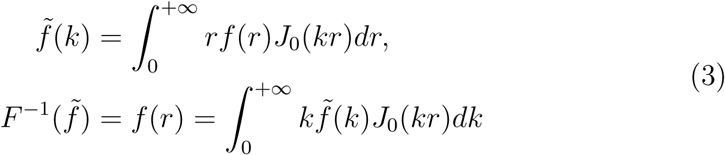

which shows that the Fourier Transform of a radial function is a radial function and vice versa.

We need to compute *f* = *d * d* where *d* is a radial function. Since

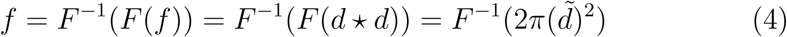

we have that *f* is a *radial* function that can be computed as

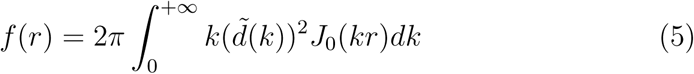

Where

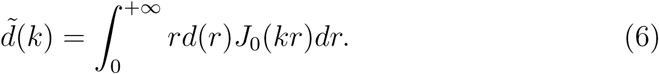

### 2. Additional Figures

**Figure 1:**
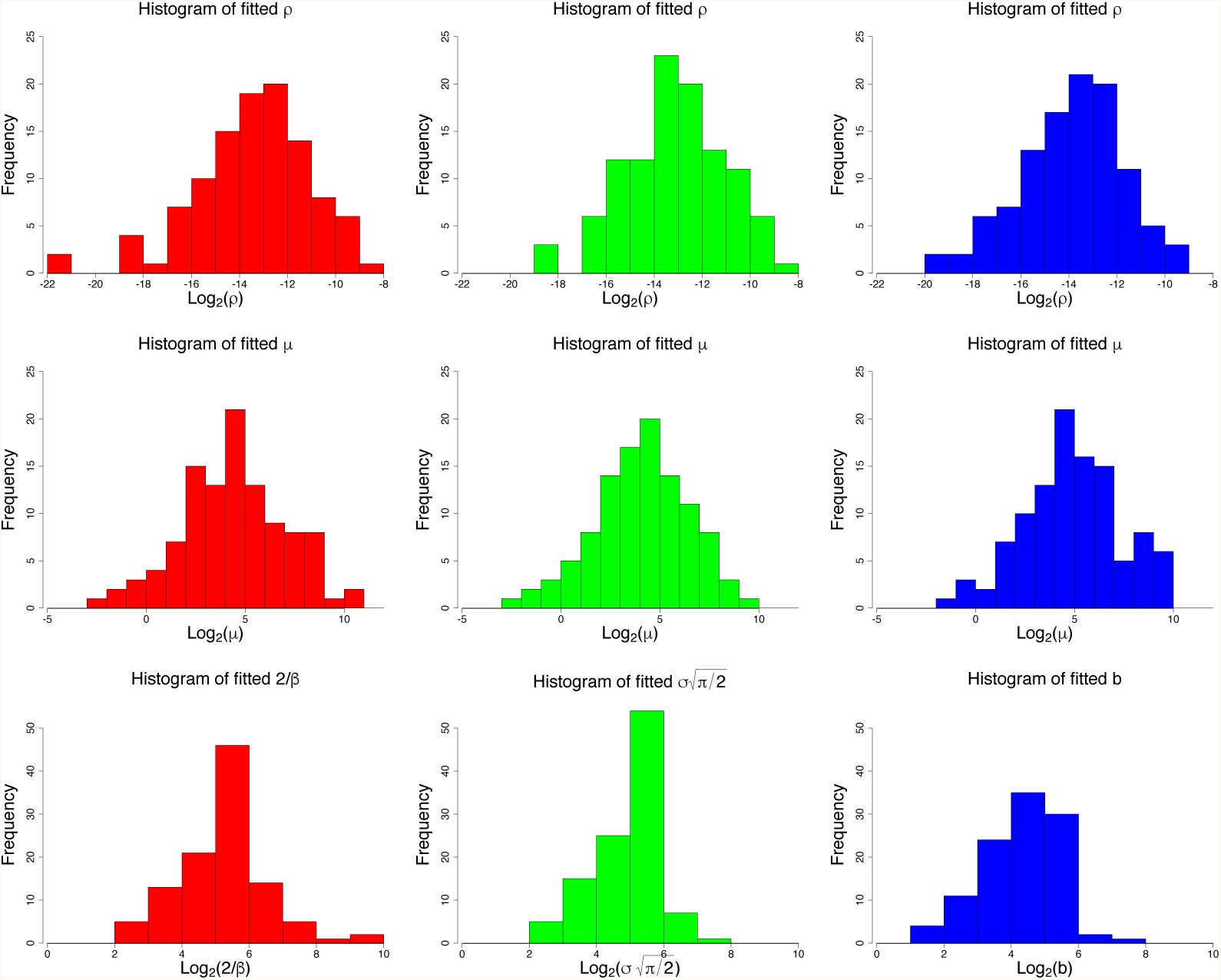
Model parameters for BCI. Fitted model parameters via the method of minimum contrast for BCI species according to the exponential process (left column), modified Thomas process (center column) and Cauchy process (right column). Form top to bottom: frequency histogram of density of cluster ρ, number of offspring per parent µ and clustering parameters β, σ and *b*, respectively. We remark that for the first two models the mean radius of cluster is well-defined and equals 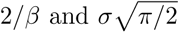 (whose histograms are shown above), while for the Cauchy process such quantity is not defined (thus above we show the histogram of log_2_ *b*).

**Figure 2:**
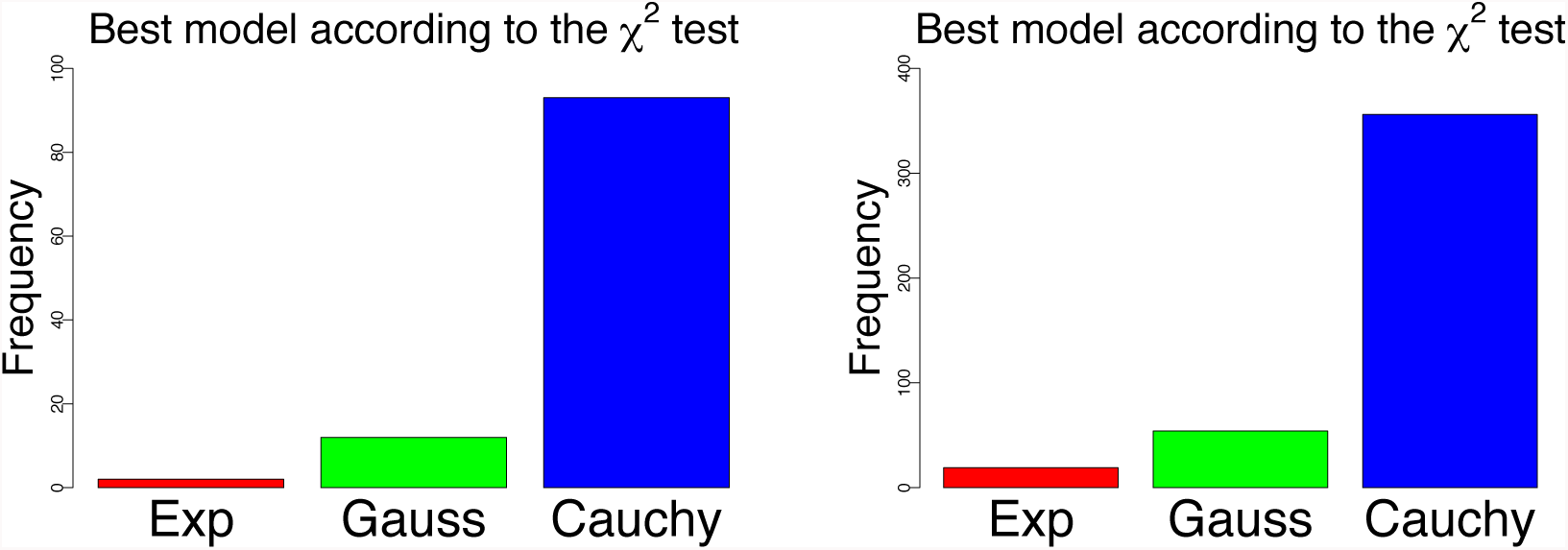
Comparison between models for BCI and Pasoh forests. We computed the goodness of each model in fitting the pair correlation function of real species through the χ^2^ test. For each species, we then select the best fitting model and we group data into frequency histograms. We find that for the overwhelming majority of the species in the BCI and Pasoh databases, the Cauchy model results to better fit the empirical pcf, followed by the modified Thomas and the exponential one.

#### 3. Relation with distance dependent Simpson index

In the paper [3] Shimatani introduces a distance-dependent Simpson index in the form

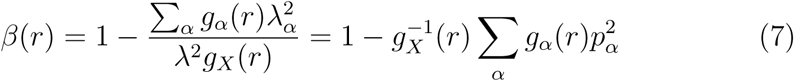

which gives the conditional probability that two individuals at distance *r* belong to different species given that there are two individuals at distance *r*.

If we set *F* (*r*) = 1 − β(*r*), the quantity *F* (*r*) is the conditional probability that two trees at distance *r* are conspecific. From the above formula one has the additional identities

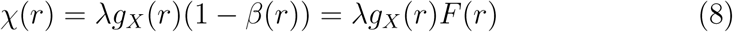

and β(*r*)*g*_*X*_(*r*) = *D*. In the paper [4] the unconditioned *F* (*r*) is analytically computed for various dispersal kernels using a neutral theory approach.

**Figure 3:**
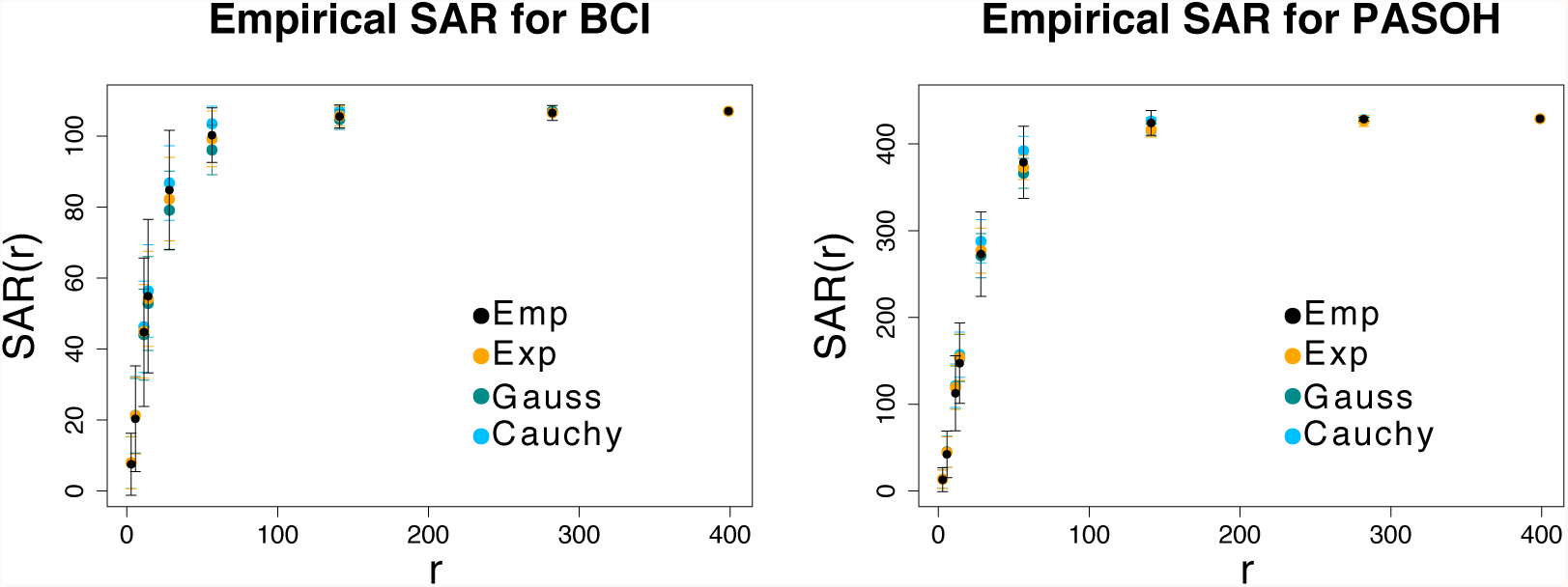
Species-Area Relationship for BCI and Pasoh. Comparison between the empirical SAR of the real BCI and Pasoh forests and the ones obtained for three artificial forests compound of species generated according to the exponential, Gaussian and Cauchy proce√sses. At each scale *r*, the 1000 × 500 window plot has been divided into *C*_*r*_ cells of side 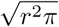. Black points and bars are mean value and three times the standard error, respectively, of the number of different species falling within each cell. Colored points and bars refer to the empirical SAR computed for three artificial forests with the same species abundance distribution as for the BCI/Pasoh but generated according to the exponential, modified Thomas and Cauchy processes. We found a good agreement between artificial and real forests, meaning that all the models are able to capture this macro-ecological pattern.

**Figure 4:**
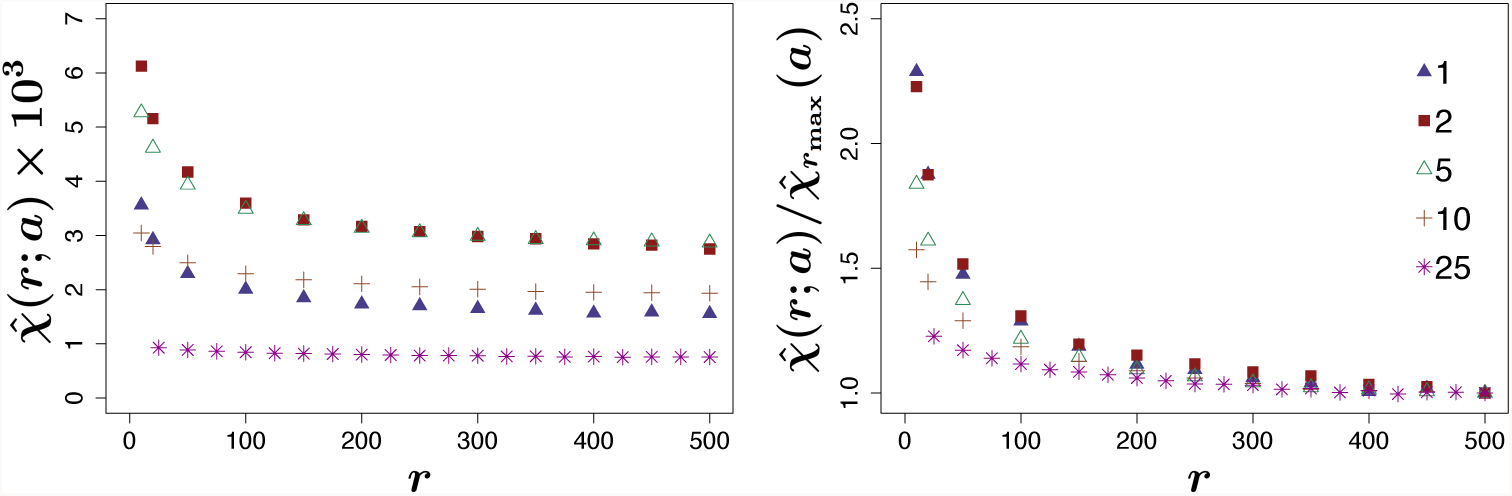
Sensitivity of Sørensen similarity index on cell size for Pasoh. We computed similarity index for Pasoh dataset considering species with more than a00 individuals. We superimposed to the 1000x500 observation window regular square grids and estimated the corresponding Sørensen index 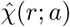 as in (22) of the main text. In figure, different colors represent different cell sizes, as in the legend. On the left panel: we can see how the choice of the cell size strongly influences the result, by ”shifting the curve 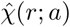 along the y-axis”. On the right panel: we divided each curve by its empirical value at the maximum considered distance, 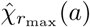 The resulting curve can be considered approximatively independent of the cell size and the error decreases with the distance.

**Figure 5:**
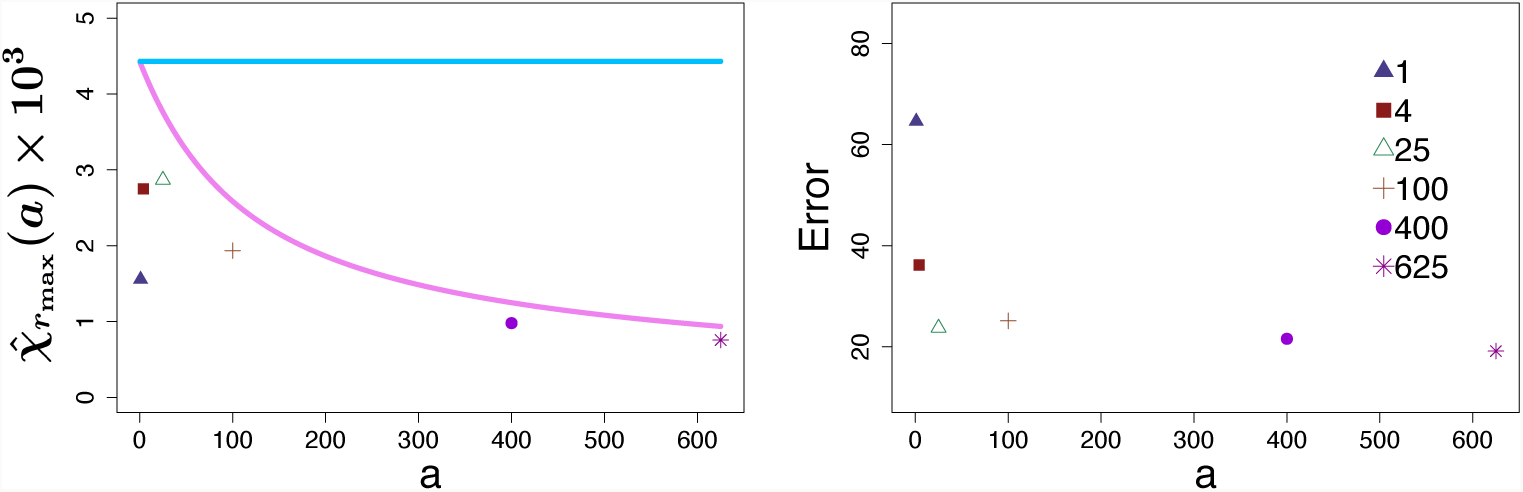
χ∞ for finite-size cells for Pasoh. On the left: the pink line represents the asymptotic value of the theoretical similarity index, χ_CSR_(*a*) as a function of the cell-area see (20) of the main text. Colored symbols are the empirical values 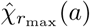 computed via (22) of the main text for cell sizes of 1, 4, 25, 100, 400 and 625 square meters, respectively, and considering only species with more than 100 individuals. The straight light blue line represents the value of 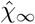 for infinitesimal cells computed through the abundances (see (18)). On the right: relative percentage error between empirical values and theoretical ones.

**Figure 6:**
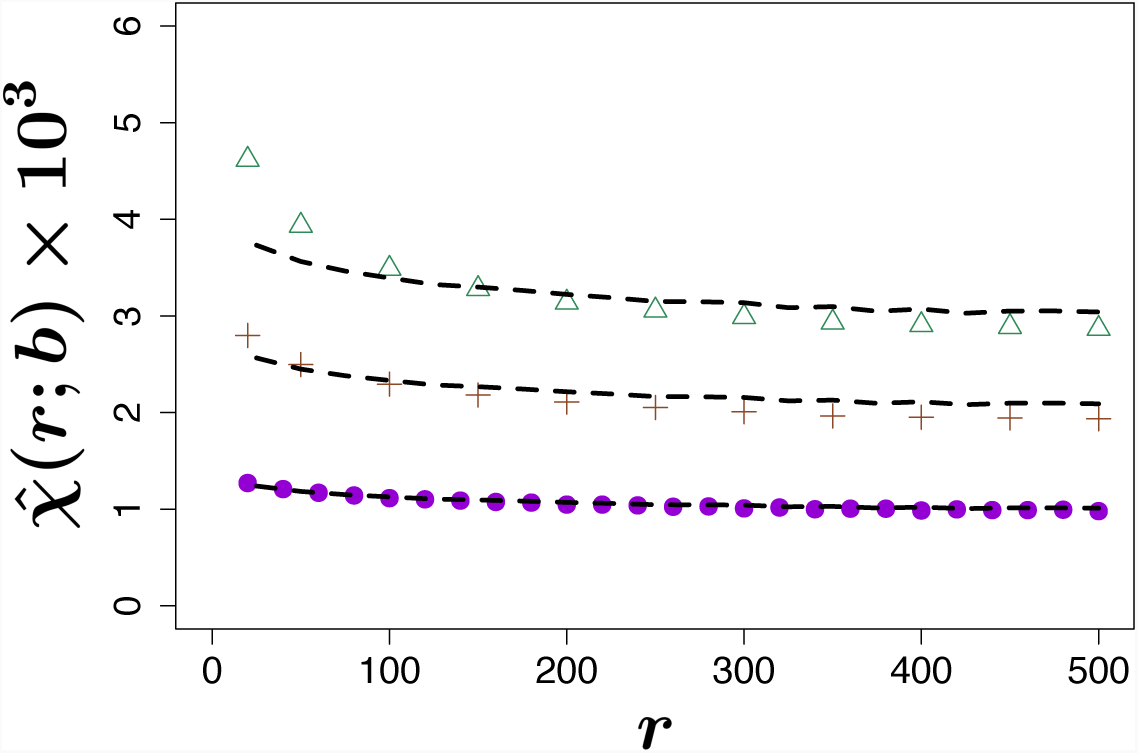
Comparison between rescaled χ estimators for Pasoh. We tested the goodness of rescaling (27) by plotting the estimator (22) for 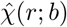 with *b* = 25 (triangles), *b* = 100 (crosses) and *b* = 400 (dots) against the rescaled one 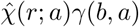 for *a* = 625 (dashed line).The difference between the two curves increases with |b a|, but there is always an excellent fit.

**Figure 7:**
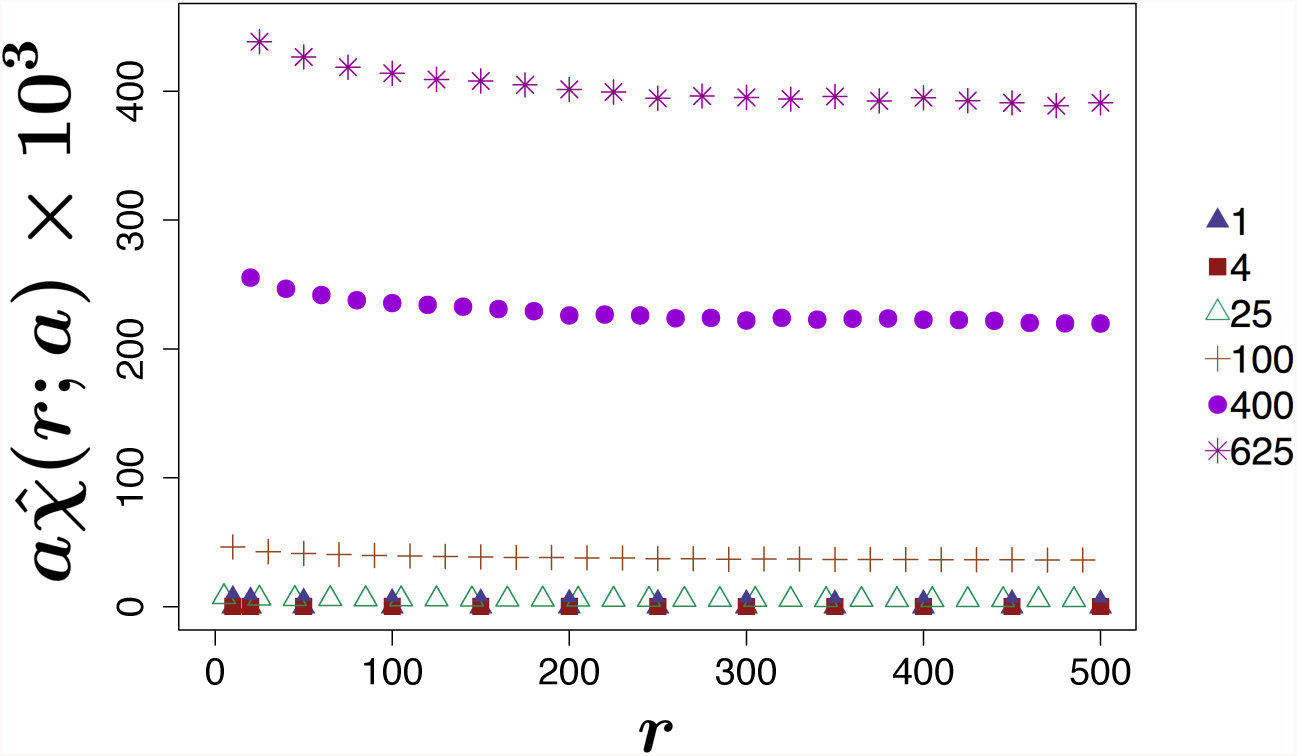
Empirical similarity index for BCI. Empirical similarity with different cell sizes. Throughout the paper we have considered the spatial density of the Sorensen similarity and its dependence on the area. In this figure, each curve 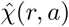 has been multiplied by the corresponding cell area *a*. The resulting curves show that even the similarity and not only its density strongly depends on the *a* parameter. Note also that the similarity curve has a higher value for larger areas, while if we consider the similarity density curves we have the opposite, similarity density is lower for bigger areas. The similarity curves on this figures show qualitatively the same dependence in the area of the curves in [2], fig.3c.

**Figure 8:**
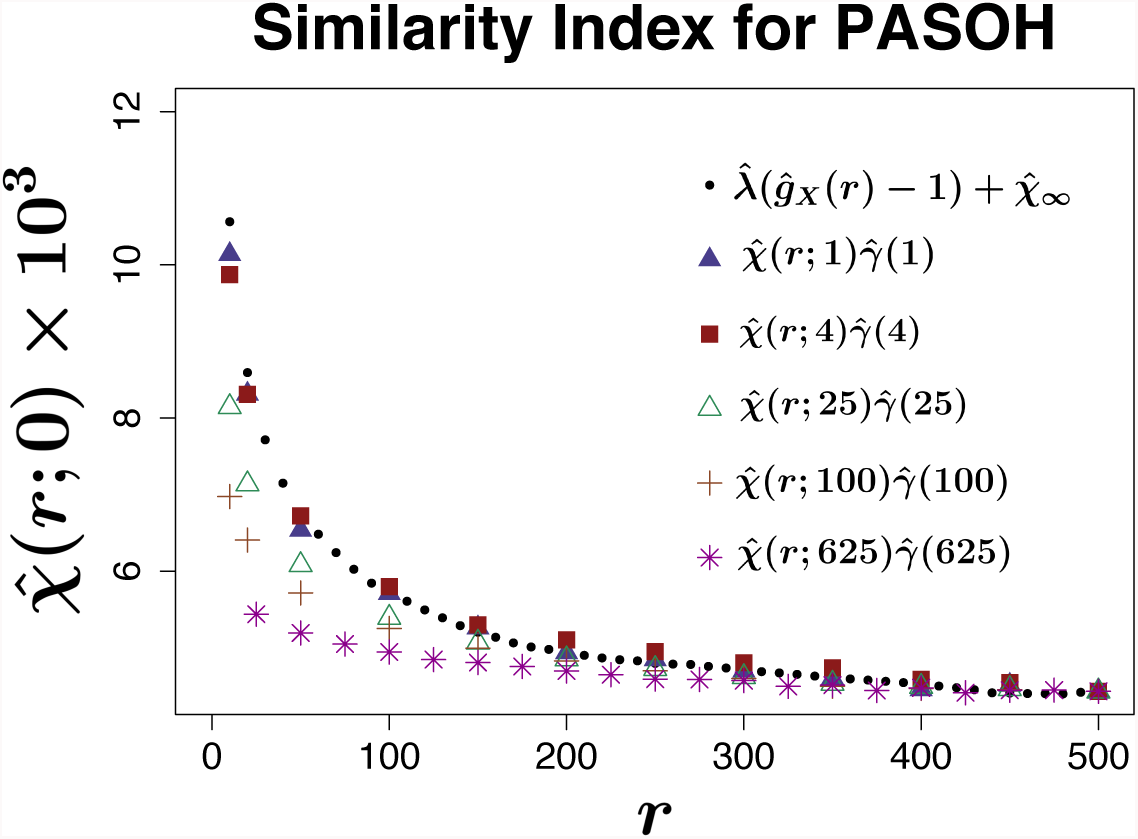
Similarity Index for Pasoh. Similarity index for Pasoh dataset computed by (26) (black dots) compared with the estimator (28) for different cell sizes a (colored symbols). The agreement between the two estimators increases as the cell size decreases. This is in accordance with the theory.

**Figure 9:**
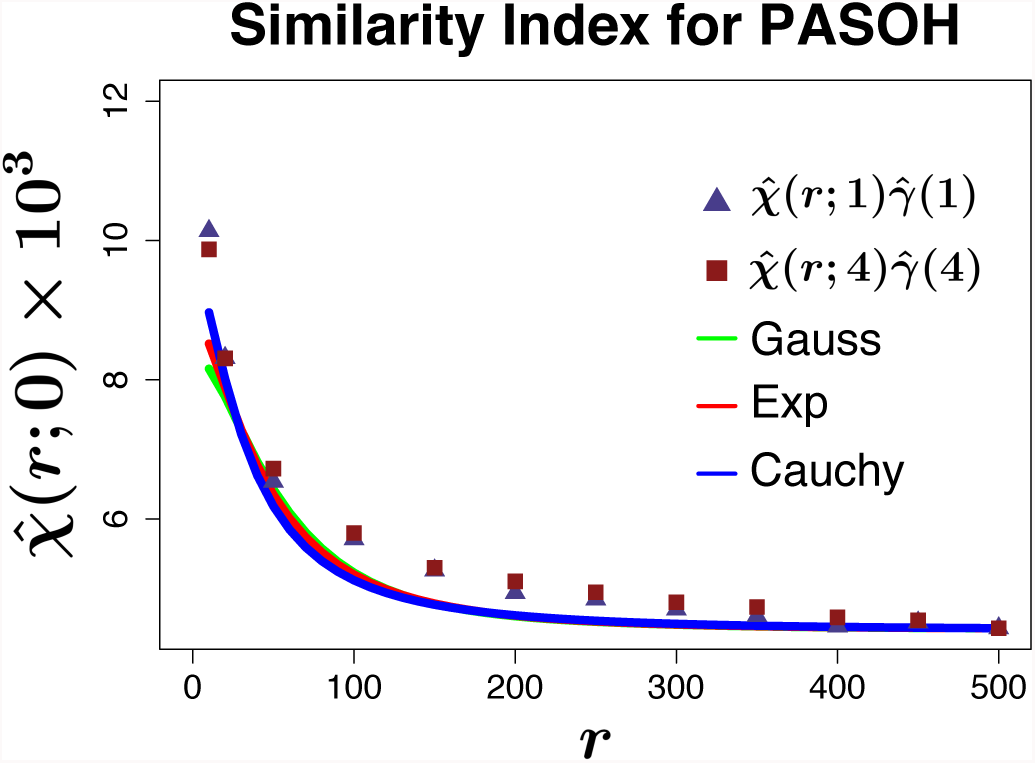
Sørensen index for Pasoh. Comparison between the empirical distancedependent Sørensen index 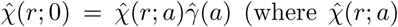 has been computed via (22) of the main text and 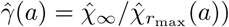 for square cells of area 1 (triangles) and 4 (squares) and the exact functional form of χ(*r*) by the three cluster models using (33) of the main text. We find a good agreement between model prediction and empirical data for all cluster type.

**Figure 10:**
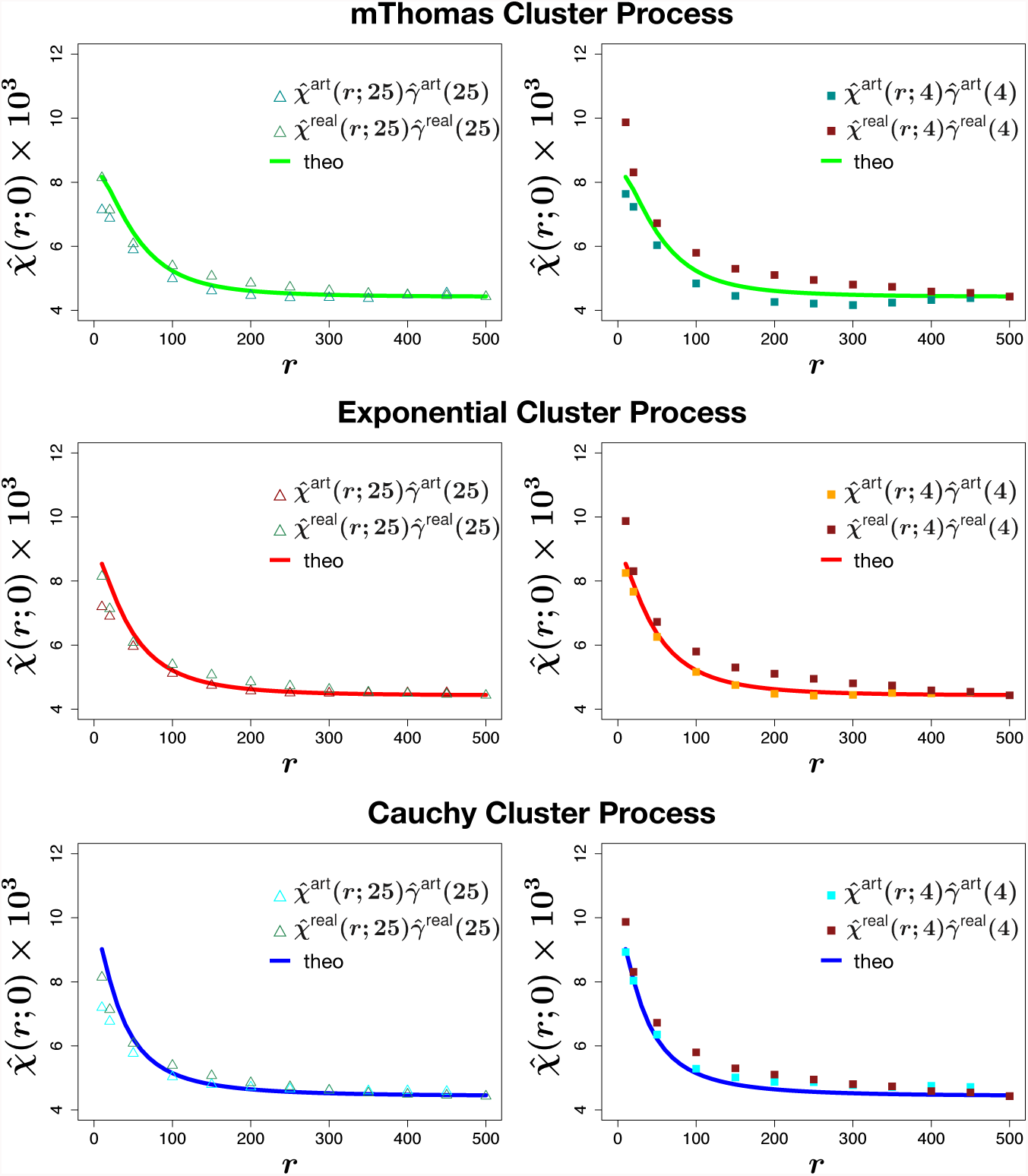
Empirical Sørensen index for Pasoh artificial forests. We generated three artificial forests as follows: for each species of Pasoh having more than 100 individuals, we generated a Poisson cluster process (modified Thomas in the top panels, Exponential in the middle panels and Cauchy in the bottom panels) within the 50 ha plot having the same number of individuals as the original species. We then computed the empirical Similarity index for new generated superposed process 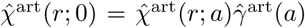 (the superscript ”art” stands for artificial forest) and compared it with the theoretical one (see (33) of the main text) and the empirical one for real Pasoh 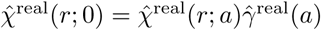 (the superscript ”real” stands for real forest), when using 25 square meters cell area (left column) and 4 square meters cell area (right column).

## Footnotes

1 We are aware that each diversity index has its own field of application and it is more or less biased. Our analytical treatment of a decay of similarity function based on a specific index will hence necessarily suffer from the same limitations of the index itself, but we are confident that our procedure can be applied to other indices.

2 If all the species have the same clustering, i.e. are described by the same pair correlation function *gα* = *g*, α *∈ {*1*, …, S}*, one has t_),_hat χ(*r*) = λ(*D −* 1)*g*(*r*), while if they are

